# Molecular Mechanism for Bacterial Degradation of Plant Hormone Auxin

**DOI:** 10.1101/2022.11.01.514793

**Authors:** Yongjian Ma, Xuzichao Li, Feng Wang, Lingling Zhang, Xing Che, Dehao Yu, Xiang Liu, Zhuang Li, Huabing Sun, Guimei Yu, Heng Zhang

**Affiliations:** Key Laboratory of Immune Microenvironment and Disease (Ministry of Education), The Province and Ministry Co-sponsored Collaborative Innovation Center for Medical Epigenetics, Haihe Laboratory of Cell Ecosystem, Department of Pharmacology, School of Basic Medical Sciences, Tianjin Medical University, Tianjin 300070, China; State Key Laboratory of Biocatalysis and Enzyme Engineering, School of Life Sciences, Hubei University, Wuhan 430062, China; YDS Pharmatech, Albany, NY, USA; The Province and Ministry Co-sponsored Collaborative Innovation Center for Medical Epigenetics, Tianjin Medical University; Tianjin Key Laboratory on Technologies Enabling Development of Clinical Therapeutics and Diagnostics, School of Pharmacy, Tianjin Medical University, Tianjin 300070, China; State Key Laboratory of Medicinal Chemical Biology, Frontiers Science Center for Cell Responses, College of Life Sciences, Nankai University, Tianjin 300071, China

## Abstract

Plant-associated bacteria play important regulatory roles in modulating plant hormone auxin levels, affecting the growth and yields of crops. A conserved auxin-degradation (adg) operon was recently identified in the *Variovorax* genomes, which is responsible for root growth inhibition (RGI) reversion, promoting rhizosphere colonization and root growth. However, the molecular mechanism underlying auxin degradation by *Variovorax* remains unclear. Here, we systematically screened *Variovorax* adg operon products and identified two proteins, AdgB and AdgI, that directly associate with auxin indole-3-acetic acid (IAA). Further biochemical and structural studies revealed that AdgB is a highly IAA-specific ABC transporter solute binding protein, likely involved in IAA uptake. AdgI interacts with AdgH to form a functional Rieske non-heme dioxygenase, which works in concert with a FMN-type reductase encoded by gene *adgJ* to transform IAA into the biologically inactive 2-oxindole-3-acetic acid (oxIAA), representing a new bacterial pathway for IAA inactivation/degradation. Importantly, incorporation of a minimum set of *adgH/I/J* genes could enable IAA degradation by *E. coli*, suggesting a promising strategy for repurposing the adg operon for IAA regulation. Together, our study identifies the key components and underlying mechanisms involved in IAA transformation by *Variovorax* and brings new insights into the bacterial turnover of plant hormones, which would provide the basis for potential applications in rhizosphere optimization and ecological agriculture.

## Introduction

Auxin is a vital phytohormone in plants, affecting almost all the aspects of plant’s life. In the meristems, auxin functions as a regulator of cell division, elongation and differentiation, determining the growth and architectures of shoots and roots^1^. Indole-3-acetic acid (IAA) is the most abundant, natively synthesized auxin in plants. IAA is not uniformly distributed but instead locally synthesized and transported directionally via the polar distributed PIN transporters in plants^2^. The resultant IAA gradients instruct plant development and are therefore dedicatedly regulated in synthesis, inactivation and degradation^3^.

Plants are associated with millions of microorganisms. The plant-microbiota interaction serves as an important layer of auxin gradient regulation, tuning plant development, phenotypes and fitness^4,5^. In particular, most rhizobacteria are found capable of producing IAA^6^, which stimulates the growth of primary and lateral roots within an optimal concentration range but causes root growth inhibition (RGI) at higher concentrations^4,7^. Apart from synthesizing IAA, a small group of plant-associated bacteria could degrade IAA as exemplified by *Pseudomonas putida* 1290, which transforms IAA to catechol using a pathway encoded by the *iac* gene cluster^8,9^. This is the currently best characterized aerobic IAA catabolism pathway and homologous *iac* loci have been identified in other IAA-degrading bacteria such as *P. phytofirmans* PsJN and *C. glathei* DSM 50014^10,11^. Additionally, an *iaa* gene cluster was found responsible for the anaerobic bacterial conversion of IAA to 2-aminobenzoyl-CoA^9^.

*Variovorax* is a gram-negative bacteria genus in the family *Comamonadaceae*, commonly found in the rhizosphere and regulating IAA levels^12^. A recent study revealed that the presence of *Variovorax* in the rhizosphere of *Arabidopsis* could completely counteract root growth inhibition (RGI) induced by a synthetic bacterial community^13^. An auxin-degradation (adg) operon is identified to be responsible for IAA degradation and thereby the reversion of RGI. The adg operon is found highly conserved among strains of *Variovorax* and unique to the *Variovorax* genus. Interestingly, the *Variovorax* auxin-degradation operon shares limited homology with the *iac* and *iaa* gene clusters, thus suggesting a different auxin IAA-degradation pathway.

In this study, we have used biochemical and structural approaches to elucidate the molecular mechanism of IAA turnover by the *Variovorax* adg operon. The two proteins, AdgB and AdgI, were found to directly associate with IAA. AdgB, featured with a two-lobed structure, was identified as an IAA-specific ATP-binding cassette (ABC) transporter solute binding protein (SBP), most likely mediating IAA uptake. Combined biochemical, structural and mass spectrometry studies revealed that IAA was transformed to the biologically inactive 2-oxindole-3-acetic acid (oxIAA) by the two-component AdgI/AdgH-AdgJ dioxygenase-reductase system in *Variovorax*. Further, we demonstrated that such an IAA-degradation property of *Variovrax* could be transplanted to *E. coli* by the transformation of a minimum gene set containing *adgH/I/J*. Together, our results uncover the major players and underlying mechanism of IAA turnover by a new auxin-degradation gene cluster in *Variovorax*, providing guidance for potential plant microbiota manipulation and optimization for ecological farming.

## Results

### AdgB directly associates with IAA

To understand the molecular mechanism of IAA turnover by *Variovorax* adg operon, we first set out to identify the components responsible for IAA binding (**Fig. 1a-b**). SBPs located in the periplasm work in concert with ABC transporters to transport a variety of compounds into bacteria^14,15^. The adg operon encodes two SBPs, AdgA and AdgB, which could potentially mediate the uptake of IAA in the rhizosphere. To determine whether AdgA and AdgB could bind with IAA, we expressed and purified the proteins for ITC measurements. As anticipated, AdgB could engage IAA with nanomolar affinity, whereas no obvious binding was detected for AdgA and IAA (**Fig. 1c and Supplementary Fig.1a**). To further understand the specificity of AdgB, we also tested the binding of AdgB with other endogenous and synthetic auxins including 4-chloroindole-3-acetic acid (4-Cl-IAA), indole-3-butyric acid (IBA), indole-3-propionic acid (IPA), 1-naphthylacetic acid (NAA) and 2,4-dichlorophenoxyacetic acid (2,4-D) (**Fig. 1b)**. All the analogs but 2,4-D could bind to AdgB albeit with substantially reduced affinities (**Fig. 1d**), indicating the specificity of AdgB for IAA. This is further supported by the results of the thermal shift assay, where IAA displayed the most significant stabilization effect on AdgB (**Fig. 1e**). By contrast, none of the auxin analogs could associate with AdgA (**Supplementary Fig. 1b**), suggesting a distinct substrate specificity for AdgA. Together, these data suggest that AdgB is an IAA-specific SBP in *Variovorax*. The high affinity of AdgB for IAA may facilitate efficient IAA uptake by *Variovorax* from the environment^16–18^.

**Figure 1.**
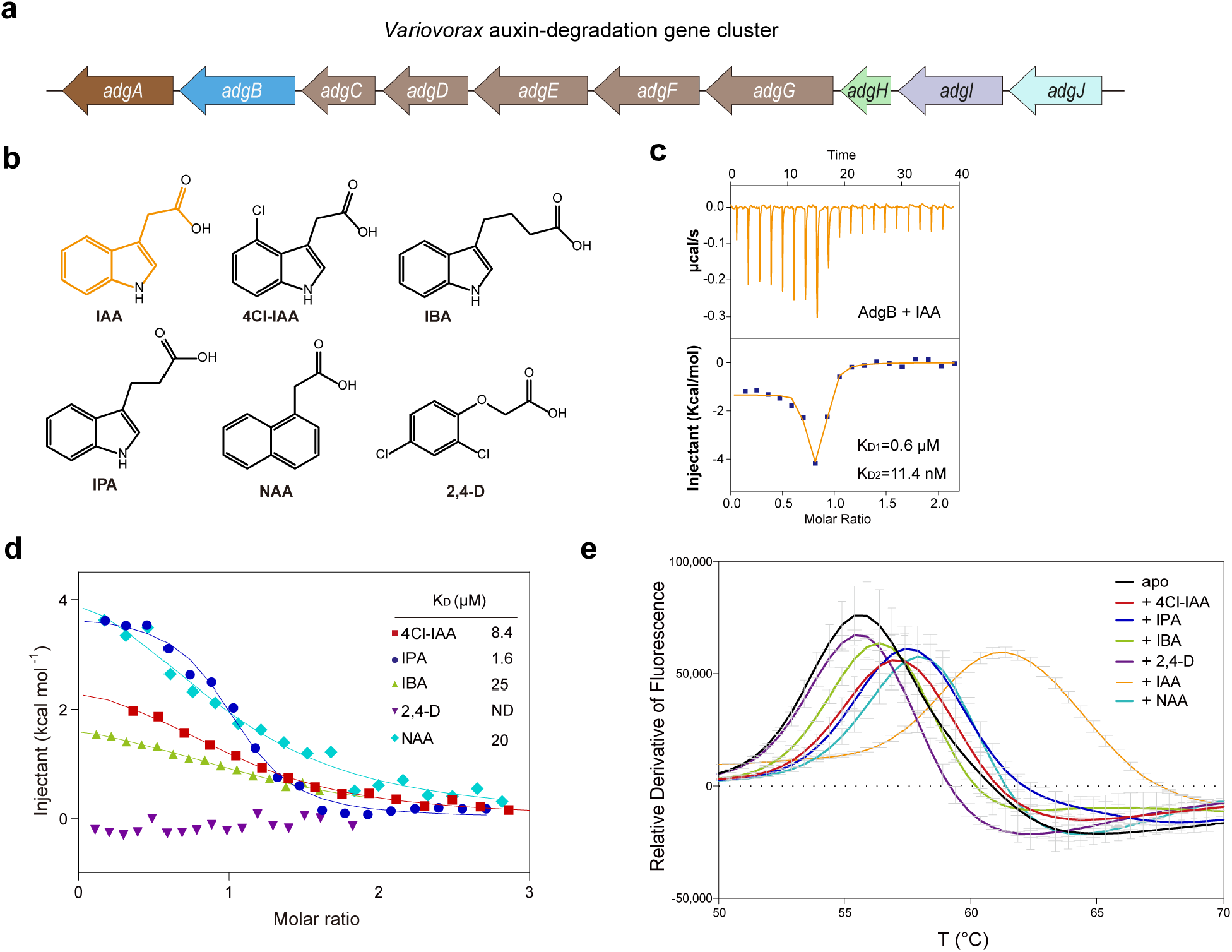
AdgB encoded by the *Variovorax* adg operon selectively binds with auxin IAA. **a**, Auxin-degradation (adg) operon in Variovorax. **b**, Chemical structures of IAA and auxin analogs. **c**, ITC measurement of IAA binding to AdgB. **d**, ITC measurements of auxin analogs to AdgB. **e**, Thermal shift assay for AdgB with IAA and analogs. Three replicates of each TSA experiments were performed.

### Structural basis for IAA engagement by AdgB

SBPs are classified into seven clusters (A-F) based on the structural similarities^15^. To further understand the IAA binding by AdgB, we next solved the crystal structure of AdgB at a resolution of 1.98 Å (**supplementary Table 1**). AdgB is composed of two rigid domains, Domain 1 and Domain 2, connected by a flexible hinge region (**Fig. 2a-b**). Both domains possess the α/β fold, consisting of a central β-sheet flanked by α-helices (**Fig. 2b**). The hinge region comprises three linkers (L1-L3) with two α-helices in linker L2, resembling SBPs in the subcluster B-III (**Fig. 2b** and **Supplementary Fig. 2a-d**), which are usually associated with type I ABC transporters for aromatic acids uptake^14,15^. Consistently, Dali search revealed a B-III SBP from *Rhodopseudomonas palustris* in complex with caffeic acid as the closest structural homology of AdgB^19^ (**Supplementary Fig. 2c**).

**Figure 2.**
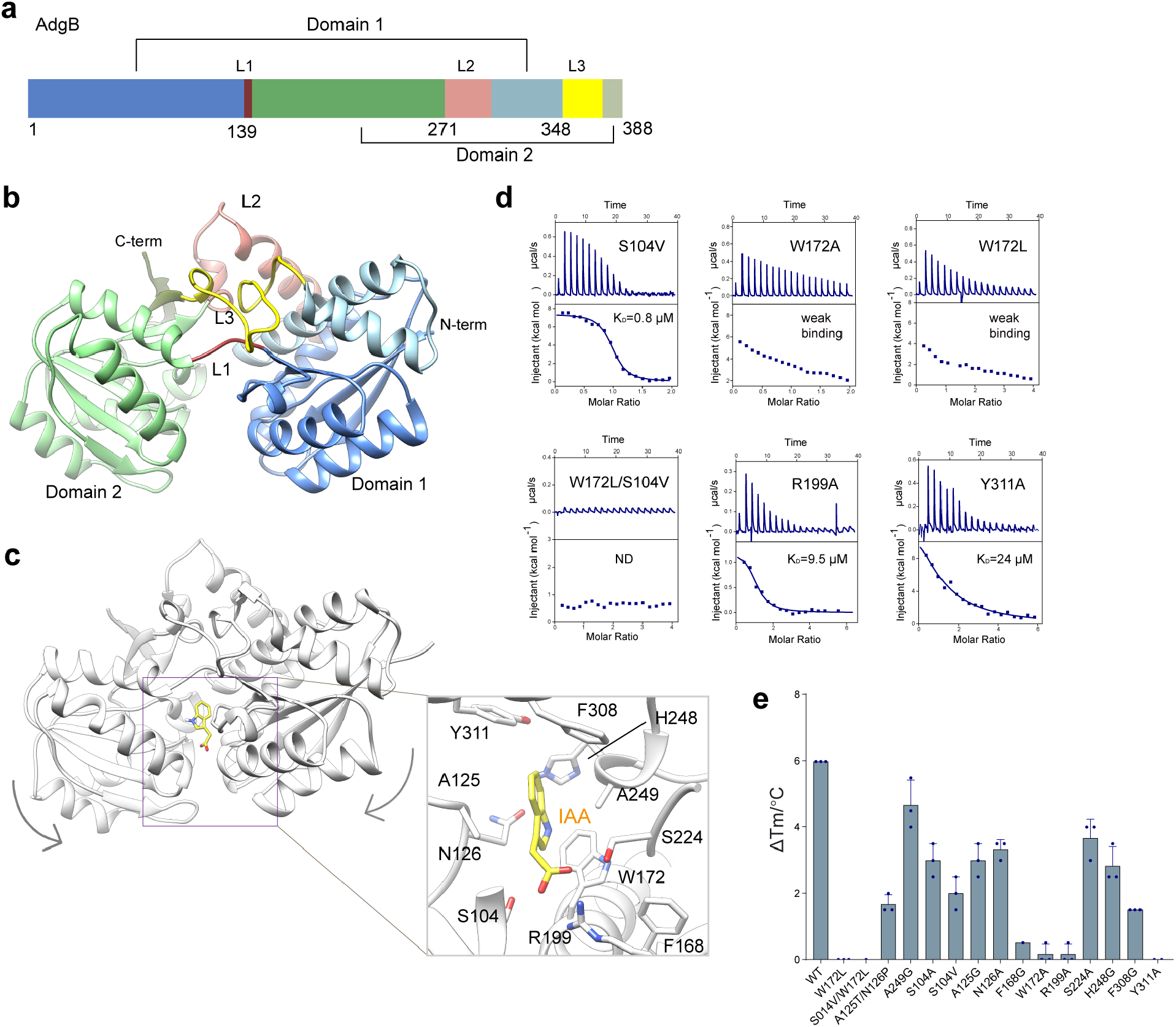
Structure of AdgB and the IAA-binding pocket. **a**, Domain organization of AdgB that is composed of two domains, Domain 1 and Domain 2. **b**, Crystal structure of AdgB. L1-L3 indicate the three linkers connecting the two lobes of AdgB. The same color scheme in **a** is applied. **c**, Molecular dynamics simulation of IAA binding to AdgB. The right panel displays the calculated IAA-binding pocket. AdgB is colored in grey and IAA is shown as yellow sticks. Key residues potentially involved in IAA binding are shown in sticks representation. **d**, ITC measurements of IAA binding to AdgB mutants. **e**, Thermal shift assay for AdgB mutants and IAA. Three replications of each TSA experiments were performed. Data are presented as mean ± s.e.m.

As co-crystallization or crystal soaking with IAA failed after extensive attempts, we therefore performed the docking and molecular dynamics (MD) simulation analysis to understand the interaction between IAA and AdgB (**Fig. 2c**). An IAA binding pocket at the interface of domain 1 and domain 2 was predicted. Notably, the computed IAA-bound structure is in a “closed” conformation in comparison to our substrate-free crystal structure (**Fig. 2b-c**), suggesting AdgB might switch from a resting “open” conformation to a “closed” conformation upon ligand binding. This is analogous to the “Venus Fly-trap”-like closure observed for other SBPs^20^. Next, mutagenesis studies were performed to validate the predicted pocket. Most tested mutations reduced or almost abolished the binding to IAA (**Fig. 2d-e**). Particularly, substitutions of Phe168, Trp172, Arg199, Phe308 and Tyr311 significantly compromised IAA binding (**Fig. 2d-e**). Phe168 and Arg199 are conserved among SBPs in the B-III cluster. Trp172 and Phe308 are moderately conserved but Tyr311 is variable among B-III SBPs (**Supplementary Fig. 2e**).

AdgA shares approximately 70% sequence identity with AdgB. To elucidate why AdgA fails to bind IAA, we predicted the structure using AlphaFold2^21^. As revealed, AdgA and AdgB share significant structural similarity (**Supplementary Fig. 2a**). Structural comparison revealed several residues located in the binding pocket in AdgB are not conserved in AdgA, such as Trp172, Ser104, Thr125 and Asn126 in AdgB (**Supplementary Fig. 2a-b, e)**. Supporting the importance of these residues in defining IAA binding, replacements with corresponding residues in AdgA obviously impaired the binding of AdgB with IAA (**Fig. 2d-e**). Notably, the double mutation replacing both Trp172 and Ser104 with those of AdgA completely abrogated the binding of IAA. Therefore, Trp172 and its neighboring residues may play important roles in determining AdgB specificity for IAA.

### Characterization of the AdgI/AdgH heterocomplex

Apart from AdgB, AdgI could also directly bind with IAA, whereas other proteins encoded by the adg operon did not (**Supplementary Fig. 3a; Supplementary Fig. 1c-d**). Sequence analysis suggested AdgH and AdgI could form a dioxygenase complex, belonging to the aromatic-ring-hydroxylating dioxygenase class. Indeed, AdgI and AdgH, analogous to the α and β subunits in canonical dioxygenase^22^, respectively, co-migrated in gel filtration assay roughly in a 1:1 molar ratio (**Supplementary Fig. 3b**). Consistently, the AdgI/AdgH complex directly binds IAA with a Kd value of about 14 μM (**Fig. 3a**).

**Figure 3.**
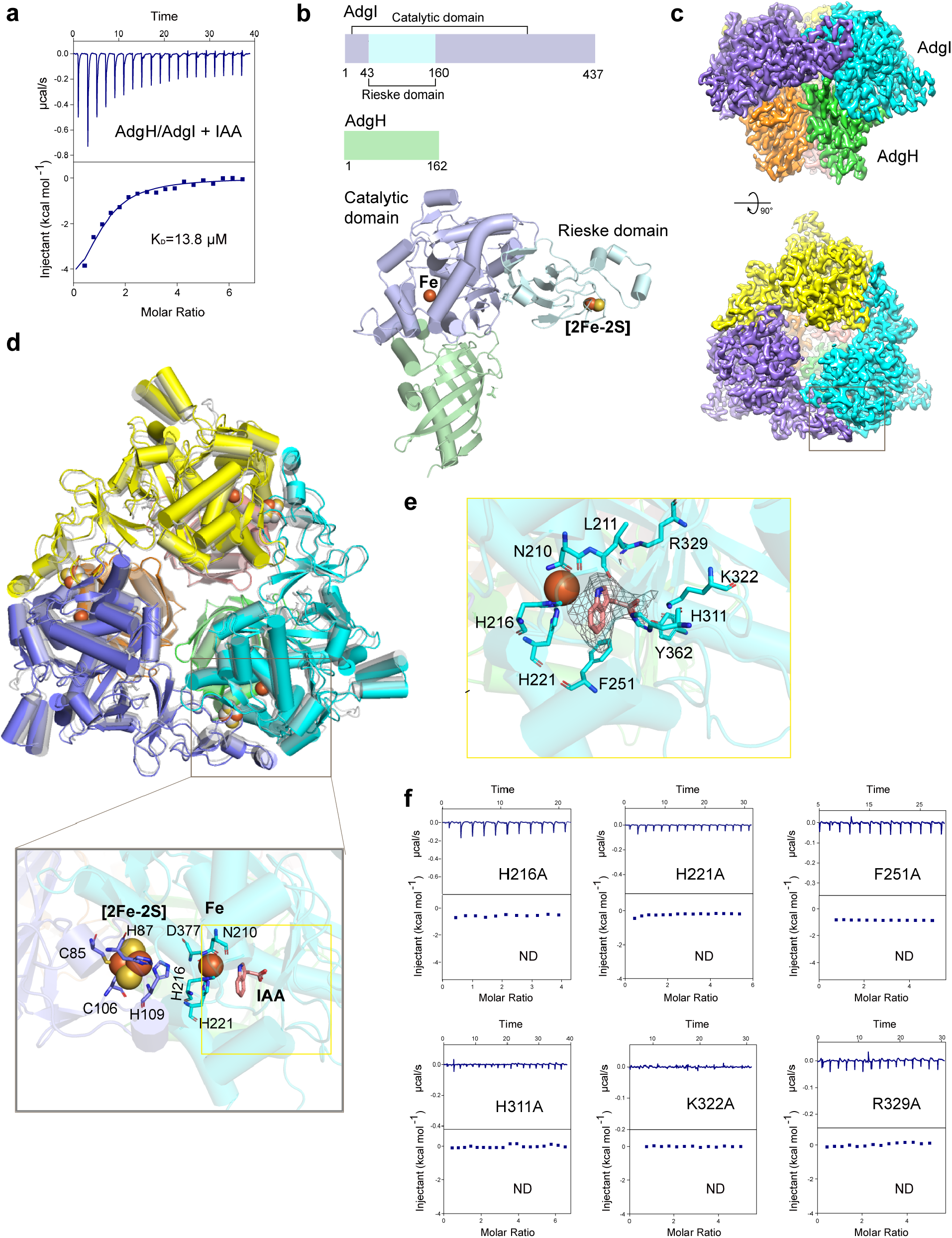
Characterization of the AdgH/AdgI Rieske dioxygenase complex. **a**, ITC measurement of IAA binding to the AdgH/AdgI complex. **b**, Crystal structure of AdgI/AdgH. Upper panel: Domain organization of AdgH and AdgI. AdgI is composed of two domains. The catalytic domain of AdgI is colored in light blue and the Rieske domain is colored in pale cyan. The AdgH is a single domain protein and is colored in pale green. **c**, 2.6 Å cryo-EM density of the AdgH/AdgI complexed with IAA. A mushroom-shaped, heterohexameric structure was resolved. The top and side views are displayed. **d**, Overlay of the apo and IAA-bound structures of AdgI/AdgH. The inset panel shows a zoom-in view of the active site composed of the [2Fe-2S] cluster, the mononuclear iron and the substrate-binding pocket. **e**, IAA binding pocket in AdgI. Key residues in the IAA binding pocket are shown as sticks. **f**, ITC measurements of IAA binding to AdgI mutants.

To further understand the structural features of the AdgI/AdgH complex, we determined the crystal structure at a resolution of 1.8 Å (**Fig. 3b and Supplementary Table 1**). Dali search revealed the biphenyl dioxygenase from *Burkholderia xenovorans* LB400 as the closest homolog^23^, which is a Rieske non-heme iron dioxygenase. The Rieske non-heme iron dioxygenases, typically composed of α and β subunits that are equivalent to AdgI and AdgH, respectively, share a conserved α3β3 configuration^24^. Both the analytical ultracentrifugation analysis and SAXS data of AdgI/AdgH supported a heterohexamer model in solution (**Supplementary Fig. 3c-d**), indicating AdgI/AdgH may share the same α3β3 configuration as other Rieske non-heme iron dioxygenases. Although one AdgI/AdgH heterodimer was observed in each asymmetric unit, the heterohexamers could be generated by crystal packing analysis (**Supplementary Fig. 4a**).

AdgI features a Rieske domain (residues 44-160) and a catalytic domain (residues 1-43 and 161-437) (**Fig. 3b**). The catalytic domain contains a core helix-grip fold with an eight-stranded antiparallel β-sheet surrounded by ten helices. The Rieske domain folds into two parallel β-sheets composed of three and four β-strands, respectively, which are connected by multiple long loops. AdgH is a single-domain protein, containing a twisted seven-stranded β-sheet and three α-helices, which may mainly serve as a scaffold protein contributing to proper assembly of the oligomeric enzyme complex.

To understand the binding of IAA with AdgI, we then determined a 2.6 Å cryo-EM structure of AdgI/AdgH complexed with IAA (**Fig. 3c, Supplementary Fig. 5 and Supplementary Table 2**). An overall mushroom-shaped heterohexamer composed of three AdgH at the “stem” and three AdgI in the “cap” was resolved (**Fig. 3c**). No profound conformational changes were observed for AdgI after binding of IAA (**Supplementary Fig. 4b**). The active site of AdgI is located at the interface of AdgI subunits, composed of a Rieske [2Fe-2S] cluster from the Rieske domain and a mononuclear iron in the catalytic domain of a neighboring subunit (**Fig. 3d**). The [2Fe-2S] is stabilized by contacts with Cys85, His87, Cys106 and His109 and the mononuclear iron coordinates with His216, His221 and Asp377. Adjacent to the mononuclear iron is the IAA binding pocket lined by polar and hydrophobic residues (**Fig. 3e)**. The resolved IAA-binding pocket of AdgI/ AdgH is largely superimposed with that of biphenyl in the biphenyl dioxygenase structure, but contains different residues to accommodate the substrate specificity (**Supplementary Fig. 4c**). As anticipated, mutation of residues in the pocket abrogated IAA binding (**Fig. 3f**). These results therefore suggest AdgI/AdgH is a Rieske non-heme dioxygenase, which may use IAA as a substrate.

### AdgI/AdgH and AdgJ reconstitute a two-component dioxygenase system for IAA degradation

The Rieske dioxygenases usually work in combination with a reductase or both reductase and ferredoxin components, forming a two- or three-component dioxygenase system, to oxidize substrates^24^. Indeed, analysis of the adg operon revealed the neighbor gene *adgJ* that encodes a reductase protein (AdgJ) (**Fig. 1a**). To test if AdgJ could work together with AdgI/AdgH to transform IAA, we purified AdgJ protein and performed the in vitro IAA degradation experiment. Indeed, IAA was transformed efficiently in the presence of both AdgI/AdgH and AdgJ, whereas neither AdgI/AdgH nor AdgJ alone could catalyze the transformation (**Fig. 4a-b**), suggesting a potential two-component dioxygenase system for IAA degradation.

**Figure 4.**
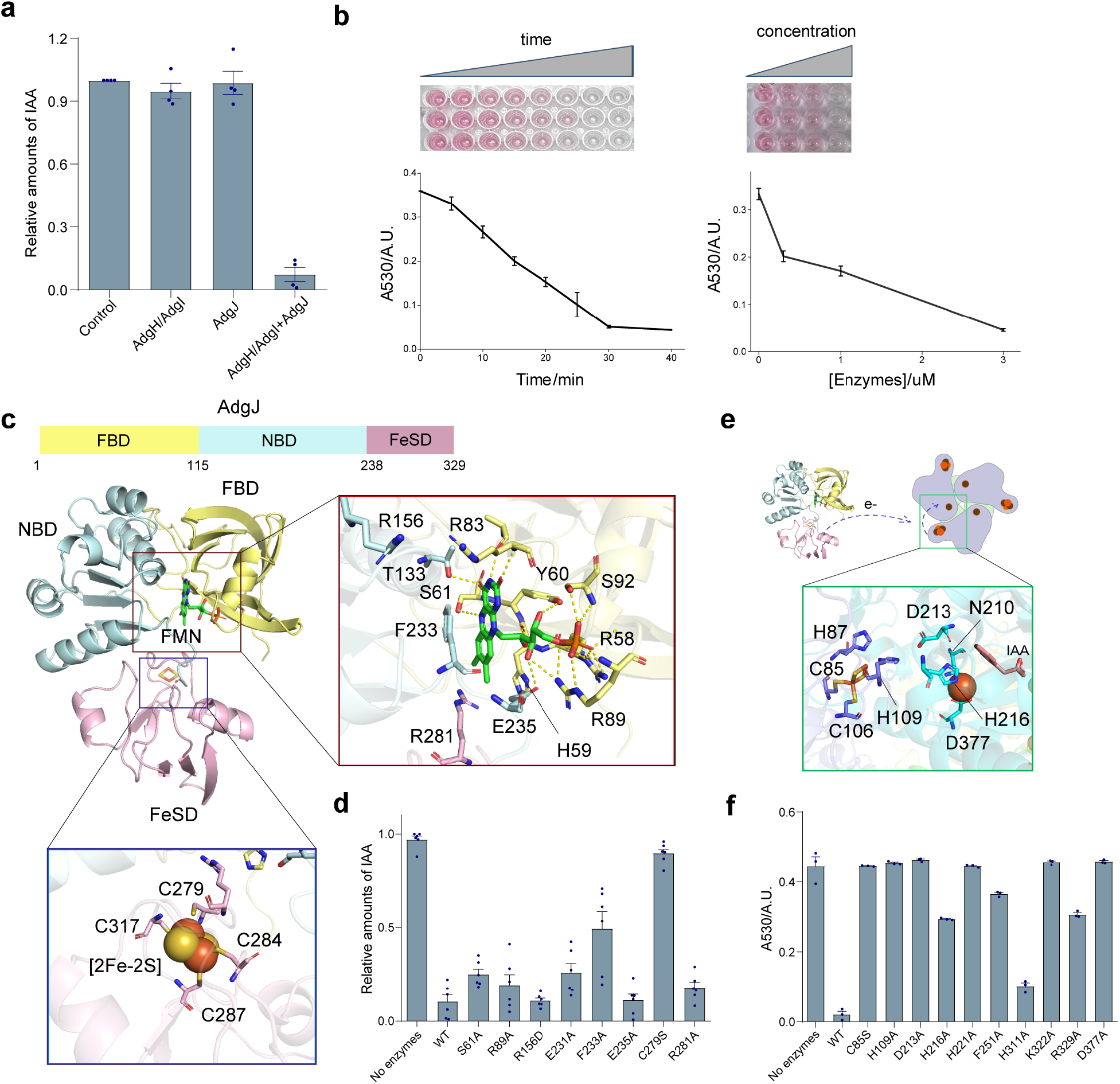
Reductase AdgJ and dioxygenase AdgI/AdgH catalyze IAA degradation. **a**, IAA degradation assay. Purified AdgI/AdgH and AdgJ were used to test the in vitro IAA degradation. The amounts of IAA were quantified using the Salkowski reagent. **b**, Time and concentration dependence of IAA degradation by AdgJ and AdgH/AdgI. **c**, Crystal structure of AdgJ. AdgJ possesses a tri-lobed structure, consisting of three domains including FBD, NBD and FeSD. The [2FE-2S] binding pocket in FeSD and the FMN binding pocket in FBD are displayed. **d**, Effects of AdgJ mutations on IAA degradation. Data are presented as mean ± s.e.m. **e**, Electron transfer from AdgJ to AdgI in catalyzing IAA degradation. The inset panel displays key residues in AdgI involved in electron transfer. **f**, Effects of AdgI mutations on IAA degradation. Data are presented as mean ± s.e.m.

Next, we determined the crystallographic structure of AdgJ at a resolution of 1.68 Å (**Fig. 4C and Supplementary Table 1**). The overall triangle-shaped protein is composed of three major domains, including an FMN-binding domain (FBD, residues 1-115), a NADH-binding domain (NBD, residues 116-238) and a plant-type [2Fe-2S] cluster domain (FeSD, residues 239-329) (**Fig. 4c**). The FBD folds into a six-stranded β-barrel and the NBD features an α/β fold, forming a five-stranded β-sheet wrapped by helices. FMN resides at the interface of the FBD and NBD domains, with the isoalloxazine ring sandwiched between the side chains of His59 and Phe233 and anchored by polar contacts with nearby residues. Further, the ribityl moieties of FMN are stabilized by polar contacts with residues Arg58, Arg89 and Ser92 (**Fig. 4c right**). FeSD is composed of a twisted β-sheet with an α-helix positioned at each side. The [2Fe-2S] cluster is coordinated by four Cys residues in two peripheral loops, facing the center of the tri-lobbed AdgJ (**Fig. 4c bottom**). AdgJ closely resembles the phthalate dioxygenase reductase (PDR) from *Pseudomonas cepacia* according to Dali analysis^25^ (**Supplementary Fig. 4d**), indicating a classical IA FMN-type reductase. However, the relative domain orientations are distinct between the two structures. While FBD and NBD are closely packed in both PDR and AdgJ structures, FeSD associates primarily with NBD in AdgJ but with FBD in the PDR (**Supplementary Fig. 4d-e**), potentially representing the different states of electron transfer^24^.

Mutagenesis studies were then performed to further confirm the functional role of AdgJ in IAA degradation. FMN is thought to mediate electron transfer from the NADH electron donor to the [2Fe-2S] electron acceptor, which further delivers the electrons to the dioxygenase for the catalysis reaction^25,26^ (**Fig. 4c, e**). As anticipated, mutation of residues involved in FMN binding compromised IAA turnover **(Fig. 4d**). Similarly, the C279S mutation impairing coordination of the [2Fe-2S] cluster significantly decreased the efficiency of the reaction. Likewise, mutation of residues involved in electron transfer and IAA binding in AdgI also significantly impaired the processing of IAA (**Fig. 4e**), supporting the critical role of the catalytic activity of AdgI in IAA degradation.

In addition to IAA, other auxin analogs also play essential roles in plant growth and development. We therefore further examined whether the AdgI/AdgH-AdgJ system could degrade IAA analogs. While 4-Cl-IAA was transformed with limited efficiency, AdgI /AdgH -AdgJ could not catalyze the turnover of IAA analogs such as IBA and IPA (**Supplementary Fig. 6a-b**). Thus, the AdgI/AdgH -AdgJ dioxygenase system appears to be selective for degradation of IAA.

### Metabolite identification of IAA degraded by the AdgI/AdgH -AdgJ dioxygenase system

Next, we set out to identify the product of IAA transformed by this new dioxygenase system. To this end, we performed the HPLC and HRMS analyses. The reacted mixture was first denatured and analyzed using HPLC. A new peak with a retention time of 5.4 min was observed accompanied with the disappearance of the IAA peak (**Fig. 5a and Supplementary Fig. 7**). Fractions from this potential product peak were then collected and subjected to the HRMS analysis (**Fig. 5b**). Two possible products were proposed based on the HRMS spectra, including the 5-hydroxy indole-3-acetic acid (5-HIAA) and the oxIAA. IAA develops the pink color with the Salkowski reagent^27^, which turned colorless after transformation by the AdgI/AdgH -AdgJ (**Fig. 4b**), indicating a product unreactive with Salkowski reagent. It therefore ruled out 5-HIAA as the potential product as that a dark gray color was observed after mixing 5-HIAA with the Salkowski reagent (**Fig. 5c and Supplementary Fig. 6**). By contrast, oxIAA was found unable to react with the Salkowski reagent (**Fig. 5c**). Furthermore, the product peak overlapped with that of oxIAA and the peak intensity increased following the addition of oxIAA to the reacted mixture in the HPLC assays (**Supplementary Fig. 7**); the adsorption spectrum of the reacted mixture also overlaid with that of oxIAA (**Fig. 5d**), both supporting oxIAA as the product.

**Figure 5.**
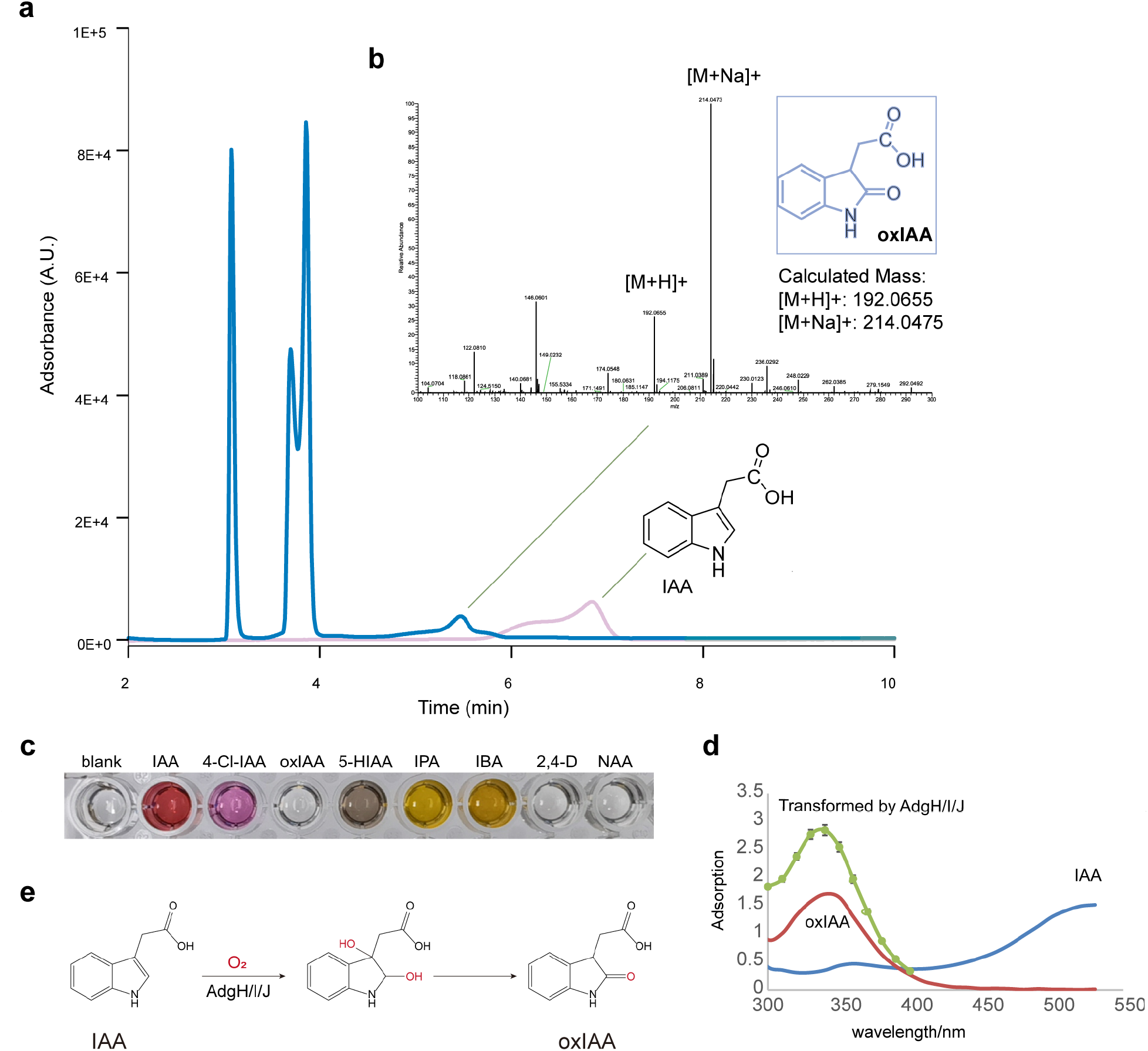
The AdgI/AdgH-AdgJ dioxygenase-reductase system converted IAA to oxIAA. **a**, HPLC analysis of IAA before and after transformation by AdgH/AdgI-AdgJ. **b**, HRMS analysis of the transformed products of IAA by AdgH/AdgI-AdgJ. The elution for potential product peak in HPLC was collected and analyzed by HRMS. **c**, Color development of IAA and analogs in Salkowski reagent. **d**, The absorbance spectra of IAA, oxIAA and the products by adgH/adgI-adgJ. **e**, The mechanism of IAA degradation by AdgI/AdgH-AdgJ. IAA was transformed to oxIAA by the incorporation of oxygen atoms from O_2_, catalyzed by AdgI/AdgH and AdgJ.

Domains of AdgI/AdgH and AdgJ would reconstitute the electron transport chain of Rieske dioxygenase, where electrons from NADH in NAD are passed to the [2Fe-2S] cluster by FMN bound to FAD of AdgJ followed by transporting to the active site of AdgI through the [2Fe-2S] clusters of AdgI/AdgH, catalyzing the incorporation of oxygen atoms of O_2_ into IAA^24–26,28^. Indeed, in the isotopic experiment with 95% H_2_O^18^ and ^16^O_2_, we found most oxIAA with m/z = 192.0655 containing ^16^O-bearing amide, indicating the oxygen was from ^16^O2 (**Supplementary Fig. 8**). Together, these data suggest that IAA was converted to oxIAA by the dioxygenase system composed of the AdgI/AdgH dioxygenase and the AdgJ reductase (**Fig. 5e**).

### Engineering of E. coli for IAA degradation

We next explored the possibility of engineering other bacterial strains for IAA degradation. Vectors containing different combinations of genes for AdgB, AdgI/AdgH and AdgJ were generated and transformed to *E. coli* to evaluate IAA degradation **(Fig. 6a)**. Consistent with the in vitro degradation assay, *adgH, adgI* and *adgJ* were required for IAA degradation by *E. coli*. Nevertheless, *adgB* appeared to be dispensable, as *E. coli* could efficiently degrade IAA regardless of the presence of *adgB* under the experimental condition (**Fig. 6b**), indicating the potential presence of endogenous IAA transporters in *E. coli*. Consistently, the loss-of-function mutations in AdgI or AdgJ abrogated IAA degradation by the engineered *E. coli*, whereas mutation of AdgB showed no obvious effect on IAA degradation (**Fig. 6c**). These results therefore suggest that incorporation of a minimum set of genes containing *adgH, adgI* and *adgJ* to *E. coli* could enable bacterial degradation of IAA.

**Figure 6.**
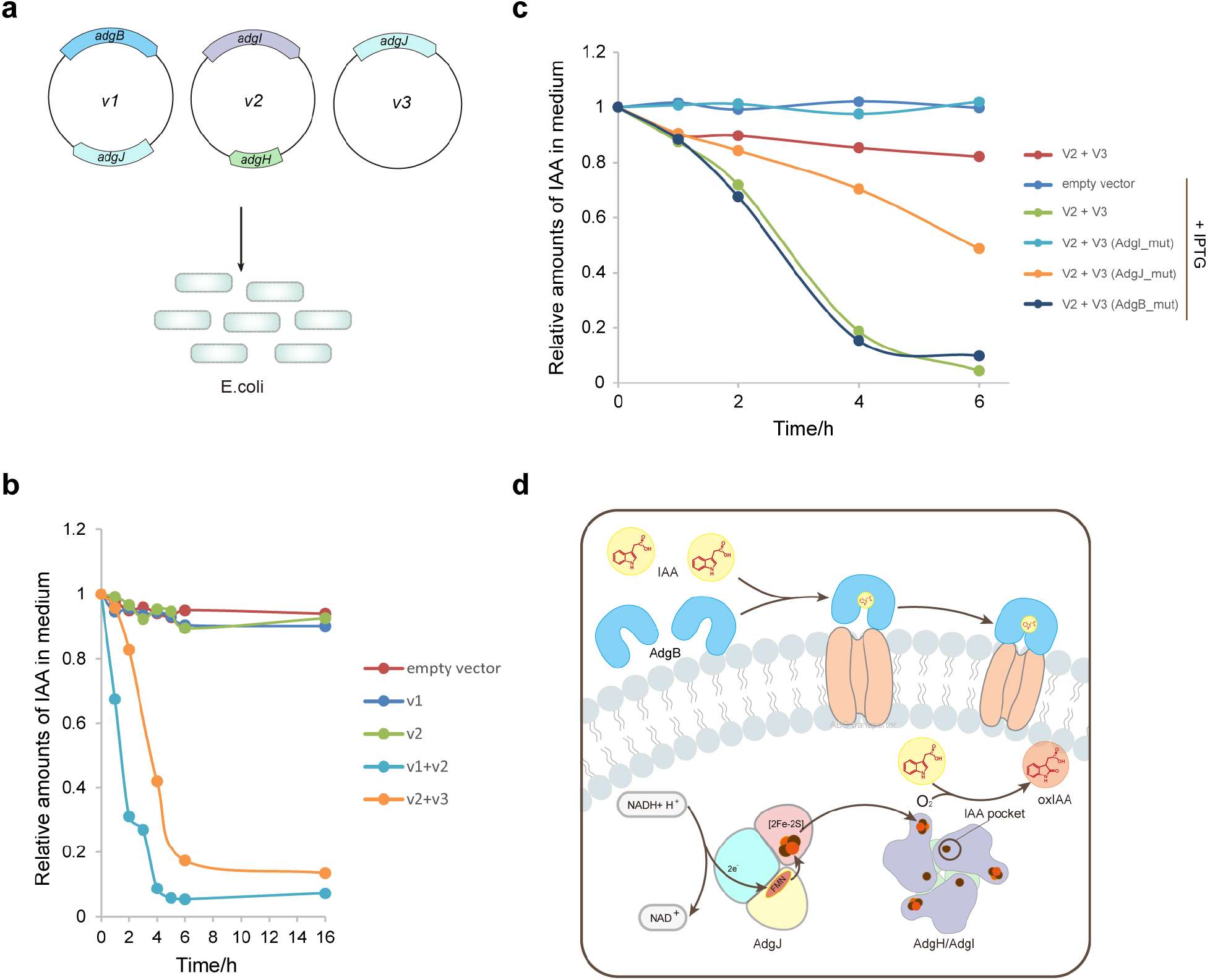
Ectopic expression of *Variovorax* genes *adgH/I/J* enables bacterial degradation of IAA by *E. coli*. **a**, A scheme for the design of the experiment for the bacterial IAA degradation assay. Vectors containing the indicated *adg* genes were transformed to *E. coli* and the degradation of IAA in the LB medium was monitored using the Salkowski reagent. **b**, In vivo IAA degradation assay for *E. coli* containing different adg genes as indicated in **a**. The presence of all three genes *adgH/I/J* resulted in efficient IAA degradation. **c**, Effects of Adg proteins mutants in the in vivo IAA degradation by *E. coli*. **d**, Schematic diagram of IAA degradation by the *Variovorax* adg operon. AdgB may mediate the efficient IAA uptake from the environment. IAA transported to the bacterial cell would be first processed by the AdgI/AdgH-AdgJ dioxygenase system and converted to the biologically inactive oxIAA.

## Discussion

Bacterial degradation of plant hormone plays an important role in regulating plant hormone levels^5,9^. Recently, a conserved adg operon coding for a new IAA degradation pathway was discovered among strains in the genus *Variovorax*, which is responsible for reversing RGI and promoting plant growth^13^. Here, we identified and characterized the proteins produced by the adg operon, directly involved in the processing of auxin IAA. Our study suggests IAA inside the bacteria could be transformed to oxIAA by an Rieske non-heme dioxygenase system constituted by AdgI/AdgH and AdgJ (**Fig. 6d**). As oxIAA is biologically inactive^3^, products of the *adgH-J* genes are likely sufficient for the bacterial downregulation of IAA in the rhizosphere to reverse RGI. Supporting our conclusion, during the preparation of this manuscript, Conway and colleagues also reported the *adgH-J* as the major genes required for IAA bacterial degradation and thereby rescuing RGI of *Arabidopsis^29^*. In addition to the AdgH-J proteins, we also found AdgB may also contribute to the IAA-degradation pathway of *Variovorax* by facilitating IAA transportation (**Fig. 6d**), although it was found dispensable for *E. coli* (**Fig. 6b**).

The adg operon encodes two SBPs that are located between the outer and inner membranes of gram-negative bacteria and typically required for nutrient and signaling molecules uptake by the ABC transporters^30^, AdgA and AdgB. Both AdgA and AdgB belong to the B-III subclass of SBPs with a binding specificity for aromatic compounds^15^. Despite the significant structural and sequence similarities, the two SBP proteins have different ligand specificity. While AdgA show no detectable binding, AdgB binds to IAA with nanomolar affinity. Considering the high binding affinity and the micromolar concentrations of IAA in the environment^16–18^, AdgB may serve as a highly efficient IAA “grasper”, working with ABC transporter in the membrane and mediating IAA uptake in *Variovorax*. This property may contribute to the supreme capability of *Variovorax* strains in degrading IAA and reverse RGI^12,13,29^. AdgB also binds with other auxins with structural similarities such as 4Cl-IAA, NAA, IPA and IBA, despite the substantially reduced affinity (μM or lower). AdgA however show no binding with these auxin analogs. Interestingly, we found AdgA could weakly associate with oxIAA. The role of AdgA in the *Variovorax* auxin-degradation pathway therefore remain to be defined.

AdgH and AdgI form a heterohexameric non-heme Rieske dioxygenase. AdgJ is an FMN-type reductase, which transfers electrons from the donor NADH to the active site of AdgI to catalyze the incorporation of oxygen atoms into IAA for the production of oxIAA. This may represent the first step in the bacterial degradation of IAA by the adg operon. The AdgI/AdgH-AdgJ dioxygenase system appears to be specific for IAA (**Supplementary Fig. 6**), similar to the ligand specificity of AdgB, indicating the preference for IAA for the adg operon pathway. The oxIAA could be further processed and decomposed by other Adg proteins in this pathway. Indeed, a recent study reports anthranilic acid as the final product of IAA degraded by the pathway encoded by the *Variovorax* adg operon^29^. Further studies are required to elucidate the functional roles of the other components in this pathway. Taken together, our studies identify the key components and elucidate the molecular mechanism underlying the most essential step in IAA degradation by the adg operon, which could be translated for applications in engineering and optimizing the rhizosphere microbiome^31,32^. For example, we have demonstrated that incorporation of a minimum gene set containing *adgH-J* could enable *E. coli* to degrade IAA.

## Methods

### Protein expression and purification

DNA sequences for genes in *Variovorax* adg operon were subcloned into expression vectors for protein purification. Genes encoding AdgA, AdgB, and AdgF were inserted into pET28-SUMO vector, and sequences for AdgC-E, AdgG-H, and AdgJ were cloned into pET28-MHL vector. Two constructs for AdgI were generated by subcloning to vectors pET28-SUMO and 13S-A (Addgene, 48323) vectors, respectively. The resultant plasmids were transformed into Rosetta DE3 cells for protein expression. AdgH and AdgI were co-expressed to obtain the functional complex. Protein expression was induced with isopropyl-β-D-thiogalactoside (IPTG) at OD600 of 0.6 and 37 °C. Cell pellets were resuspended in the binding buffer (25 mM Tris-HCl pH 7.5, 500 mM NaCl, 5 mM imidazole and 2 mM β-mercaptoethanol) and broken by sonication. Ni-NTA resins (Qiagen) were added and incubated with the cleared lysate at 4 °C for 1 h before being extensively washed with the washing buffer (binding buffer supplemented with 15 mM imidazole). Target protein was eluted with the elution buffer containing 300 mM imidazole. The expression tags were removed by TEV protease. Tag-free proteins were concentrated and loaded onto the HiTrap SP column (Cytiva) for further purification. Gel-filtration was performed at the last step of purification on a ÄKTA Pure system with running buffer containing 25 mM Tris-HCl pH 7.5, 150 mM NaCl, 2 mM dithiothreitol (DTT). Peak fractions were collected and concentrated for use. Mutant proteins were purified in the same way as described above.

### Isothermal titration calorimetry (ITC)

ITC experiments were performed using a MicroCal PEAQ-ITC system (Malvern). All the experiments were performed at 25 °C in a buffer containing 25 mM Tris-HCl pH 7.5 and 150 mM NaCl (unless otherwise stated). A typical titration experiment involved 18 injections of IAA solution (500-2000 μM) into the cell containing protein of interest (30–100 μM) with a time spacing of 120s. Data was analyzed using the MicroCal PEAQ-ITC analysis software. All the measurements were repeated at least two times.

### Thermal shift assay (TSA)

TSA was performed to examine the binding of Adg proteins with IAA and analogs. Ligands concentrations were tested at 2-100-fold excess (10-1000 μM) relative to the proteins (5 μM). The fluorescence dye SYPRO Orange at a final of 200-fold dilution was used to monitor the unfolding of proteins. The protein and ligand were incubated for 10 min at room temperature before the addition of SYPRO Orange dye. The QuantStudio™ 1 Plus real-time PCR system (Thermo Fisher) was used to control the temperature increasing from 25 °C to 99 °C with an increment of 0.5 °C and to monitor the fluorescence (excitation: 520 ± 10 nm and emission: 558 ± 10 nm). Each tested group was run in triplicate and repeated. Data was imported to the Protein Thermal Shift™ software for analysis and plotting.

### Gel-filtration assay

The gel-filtration assay for the AdgI/AdgH complex was carried out using the HiLoad 200 column (Cytiva). AdgI/AdgH purified by the cation exchange chromatography was concentrated to 20 mg/ml and loaded into the column pre-equilibrated with the running buffer containing 25 mM Tris-HCl pH 7.5, 150 mM NaCl and 2 mM DTT. The peak fractions were collected and analyzed with SDS-PAGE.

### Analytical Ultracentrifugation analysis

The purified AdgI/AdgH complex sample was diluted to 0.5 mg/ml using the sample buffer containing 25 mM Tris-HCl pH 7.5 and 150 mM NaCl and subjected to the sedimentation velocity measurements using an Optima XL-I analytical ultracentrifuge (Beckman-Coulter). Data was analyzed with the SEDFIT and SEDPHAT programs^33,34^.

### Small-angle X-ray scattering

The AdgI/AdgH complex was concentrated to 20 mg/ml in a buffer containing 25 mM Tris-HCl pH 7.5, 280 mM NaCl and 2 mM DTT. Small-angle X-ray scattering (SAXS) data was collected on beamline BL19U2 at Shanghai Synchrotron Radiation Facility (SSRF) with a Pilatus detector. One-dimensional intensity curves were obtained by the ATSAS package^35,36^. SAXS data processing and analysis were carried out using PRIMUS software within the ATSAS package^37^. Distance distribution and molecular weight were obtained from GNOM^38^. Ab initio modeling and low-resolution 3D shape envelopes reconstruction were performed by DAMMIF^39^. CRYSOL and SREFLEX were used to test the fit quality of reconstructed models with the experimental SAXS profiles^36^.

### In vitro IAA and analogs degradation assay

The in vitro degradation assay was performed to test the capability of the AdgI/AdgH-AdgJ dioxygenase system in transforming IAA and analogs. The reaction mixture was composed of 100 mM Tris-HCl pH7.5, 250 μM reduced NADH, 10 μM ammonium iron (II) sulfate, and 3 μM of each enzyme component. The reaction was initiated by the addition of substrate (0.1-0.5 mM) at room temperature (unless otherwise stated). The mixture was vortexed periodically to introduce oxygen for the reaction and was boiled to stop the reaction. After pelleting the precipitants, the supernatant was collected and mixed with two volumes of Salkowski reagent made of 0.5 M ferric chloride and 35% perchloric acid for the quantification of residual substrate.

### HPLC and HRMS analysis of IAA metabolite

The reacted mixture from the IAA degradation assay was denatured by heat and the supernatant was collected for high-performance liquid chromatography (HPLC) analysis on a Shimadzu LC-40 HPLC machine equipped with a InertSustain 5 μm C18 HPLC column (4.6 × 250 mm). The UV detection was carried out at 260 nm, following separation using water (solvent A) and methanol (solvent B) as mobile phase with a linear gradient elution at a flow rate of 0.8 ml/min as follows: 0-5 min, 5% B; 5-15 min, 10% B; 15-20 min, 50% B; 20-22 min, 50% B; 22-25 min, 95% B; 25-33 min, 95% B. The peak fraction with a retention time of about 5.4 min was collected and re-dissolved in water containing 10% methanol after removing the solvent under vacuum. High-resolution mass spectrometry (HRMS) was performed with an LTQ-Orbitrap mass spectrometer (Thermo Fisher). The isotopic experiment was carried out in the solution containing 95% H_2_O^18^ as well as ^16^O_2_ in the air following the same protocol as described above.

### IAA bacterial degradation assay

*Variovrax adg* genes *adgB, adgH-I* were transformed to *E. coli* to screen for the minimum set required for IAA degradation. The *E. coli* cells were cultured in the LB medium till OD600 of 0.6 at 37 °C and IPTG was then added to induce Adg proteins expression. Two hours later, 0.2 mM IAA was added to the medium and the cells were grown for 16 hours, during which culture aliquots were collected and the amounts of IAA were determined using the Salkowski reagent.

### Crystallization, data collection and structure determination

Purified AdgB, AdgI/AdgH complex, and AdgJ were concentrated to 15-20 mg/ml. Protein crystallization was performed by mixing 1 μl of protein solution with 1 μl of the crystallization buffer using the sitting drop method. Crystals of AdgB protein were grown in a reservoir solution containing 27% w/v Polyethylene glycol 3,350, 0.1 M Tris pH 8.60.1 M cobalt (II) chloride hexahydrate. AdgI/AdgH crystals were obtained at 18 °C in a reservoir solution containing 28% (v/v) 2-methyl-2,4-pentanediol, 0.05 M sodium cacodylate trihydrate pH 6.0, 0.05 M magnesium acetate tetrahydrate, 40% (v/v) 1,3-propanediol. Crystals of AdgJ protein were grown at 18 °C in a reservoir solution containing 0.1 M Bis-tris propane pH 7.1, 1.2 M sodium malonate pH 7.0, 0.1 M calcium chloride dihydrate. Crystals were cryo-protected in the mother liquor supplemented with 20% glycerol before flash-frozen in liquid nitrogen for data collection.

X-ray diffraction datasets were collected at beamline BL19U1 of Shanghai Synchrotron Radiation Facility (SSRF). Structure determination was performed by CCP4I2 package^40^. Structure of AdgB was determined by molecular replacement using a predicted model from Alphafold2^21^. Structure of the AdgI/AdgH protein complex was solved by molecular replacement using the crystal structure of toluene 2,3-dioxygenase (PDB ID:3EN1). Initial model improvement was performed manually in COOT^41^. Model refinement was performed by phenix.refine^42,43^ and CCP4I2 package^40^. Data processing and structural refinement statistics are shown in the **Supplementary Table 1**.

### Docking and molecular dynamics (MD) simulation

Docking and molecular dynamics simulation analysis were performed to define the binding site of IAA in AdgB. The initial binding pocket was selected by referring to the homologous structure (PDB ID: 4FB4). Molecular docking of IAA anion to the potential binding site was performed using AutoDock Vina^44^. The revealed initial binding pose served as the start point for MD simulation. MD simulations were performed using the OpenMM and Ambertools software packages^45,46^. The anion IAA was modeled using the generalized Amber force field, and the protein structures were modeled using the Amber14SB force field^47^. The protein-ligand complexes were solvated with TIP3P water molecules in a cubic box with a length of 12.0 Å, and sodium or chloride ions were added to neutralize the systems. For all the simulations, all the bonds that involved hydrogen atoms were constrained and the Langevin dynamics thermostat was applied to keep the temperature at 300 K.

### Cryo-EM sample preparation, data collection and structure determination

Purified AdgI/AdgH complex was incubated with 20-fold molar excess IAA. Aliquots of 4 μL samples were applied to glow-discharged holey carbon girds (Cu, R1.2/1.3, 300 mesh, Quantifoil). The grids were blotted with force 2 for 3 seconds and plunged into liquid ethane using a Vitrobot (FEI Thermo Fisher). Cryo-EM data were collected with a Titan Krios microscope (FEI) operated at 300 kV and images were collected using EPU at a nominal magnification of 105,000x (resulting in a calibrated physical pixel size of 0.85 Å/pixel) with a defocus range from −1.2 um to −2.2 μm. The images were recorded on a K3 summit electron direct detector (Gatan). A dose rate of 15 electrons per pixel per second and an exposure time of 2.5 seconds were used, generating 40 movie frames with a total dose of ~ 54 electrons per Å2. A total of 2122 movie stacks were collected (**Supplementary Table 2**).

The movie frames were imported to RELION-3^48^. Movie frames were aligned using MotionCor2^49^ with a binning factor of 2. Contrast transfer function (CTF) parameters were estimated using Gctf^50^. Particles were auto-picked and extracted from the dose weighted micrographs. The following steps, including 2D classification, generation of initial model, 3D classification and 3D refinement, were performed in cryoSPARC^51^. 1,717,755 particles were selected after 2D classification for further processing. Heterogeneous refinement was performed with the three initial models to distinguish different conformational states. 3D refinement was performed with C1 symmetry for the selected class representing the intact complex. After confirming that the IAA molecule is present in all the copies of the AdgI/AdgH complex. A second round of 3D refinement with C3 symmetry was performed, converging at a resolution of 2.59 Å.

The map was sharpened using Phenix Autosharpen tool^52–54^. The apo AdgI/AdgH crystallographic structure (**Supplementary Table 1**) was used as the initial template. Manual adjustments and refinements were performed with Coot^41^. Iterative real-space refinement was then performed in Phenix using phenix.real_space_refine^53^. The quality of the model was analyzed with MolProbity in Phenix^55^. Refinement statistics are summarized in **Supplementary Table 2**.

## Data Availability

The atomic coordinates of the structures have been deposited in the Protein Data Bank under accession codes 7YLT (AdgB), 7YLS (AdgI/AdgH), 7YLR (AdgJ) and 8H2T (AdgI/AdgH-IAA). The cryo-EM map of AdgI/AdgH in complex with IAA has been deposited to the Electron Microscopy Data Bank under the corresponding accession code EMD-34443.

## Acknowledgements

This work was supported by the National Natural Science Foundation of China (32071218 to H.Z.; 32200496 to G.Y.; 22007072 to H.S.). We thank the staffs from the BL17B/BL18U1/BL19U1/BL19U2/BL01B beamlines of the National Facility for Protein Science (NFPS) at Shanghai Synchrotron Radiation Facility for assistance in crystallographic data collection. The cryo-EM data was collected using the Cryo-Electron Microscopy Facility of Hubei University.

## Author Contributions

H.Z., G.Y. and H.S. conceived the study. Y.M. and L.Z. performed the molecular cloning, protein purification and biochemical assays with the help from G.Y. and H.Z. Y.M performed the protein crystallization and X.L. processed the crystallographic data and built the atomic models. Z.L. and F.W. collected and analyzed the cryo-EM data. G.Y. built and refined the structure model. H.S. and D.Y. performed the HPLC and HRMS analyses. X.C. provided the docking and molecular dynamics simulation analysis of the structure. X.L. performed the analytical ultracentrifugation experiment. G.Y. wrote the manuscript with contributions from H.Z..

---

**Supplementary Figure 1.**
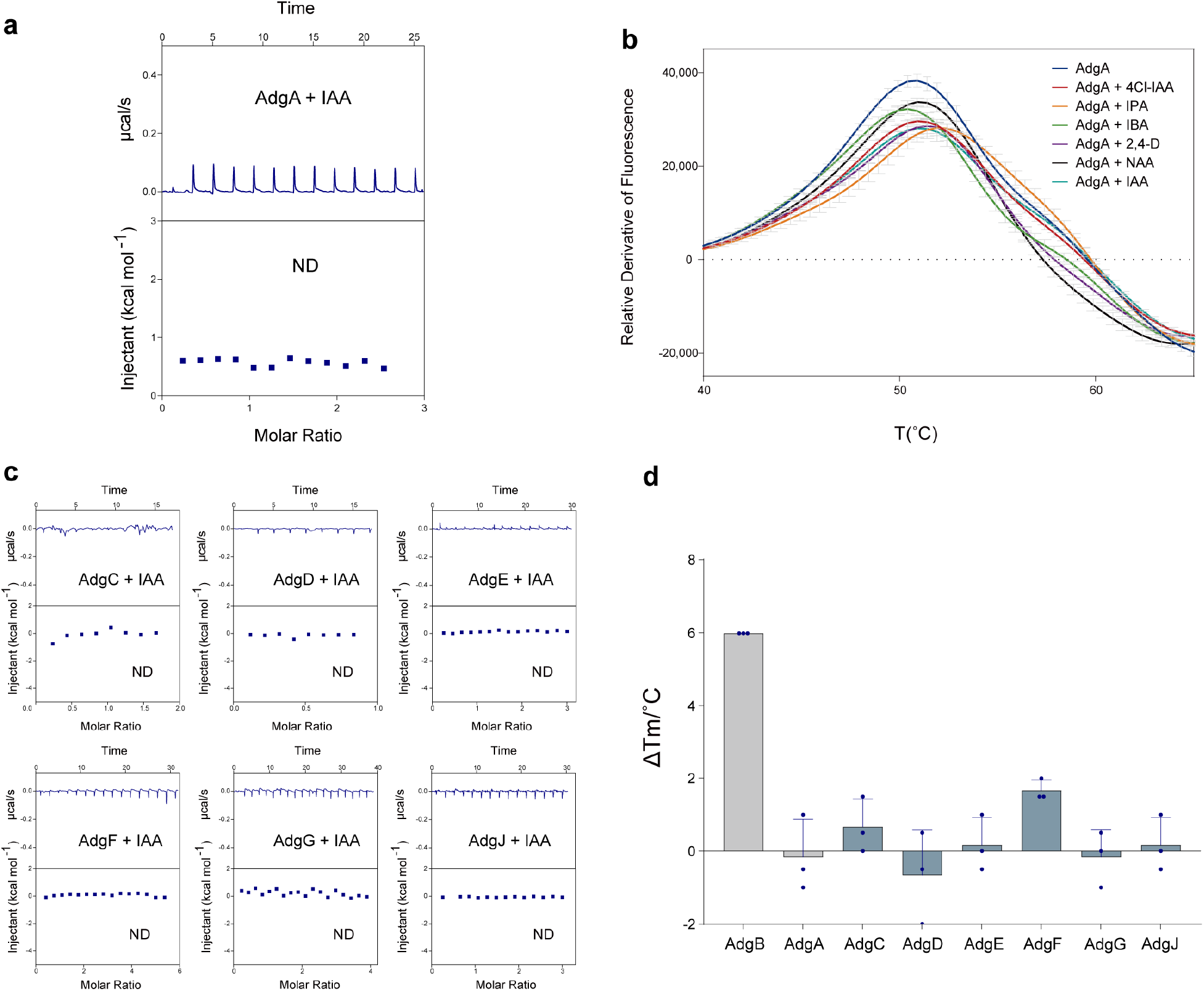
Interaction between Adg proteins and IAA. **a**, ITC measurement of IAA to the AdgA SBP. **b**, Thermal shift assay for AdgA with IAA and different analogs including 4Cl-IAA, IPA, IBA, 2,4-D and NAA. Three replicates of each TSA experiments were performed. **c**, Integrated heat plots for ITC measurements of Adg proteins. Measurements for proteins AdgC-G and AdgJ with IAA are displayed. ND: no binding detected. **d**, Tm changes for Adg proteins and IAA in the thermal shift assay. Three replications of each TSA experiments were performed.

**Supplementary Figure 2.**
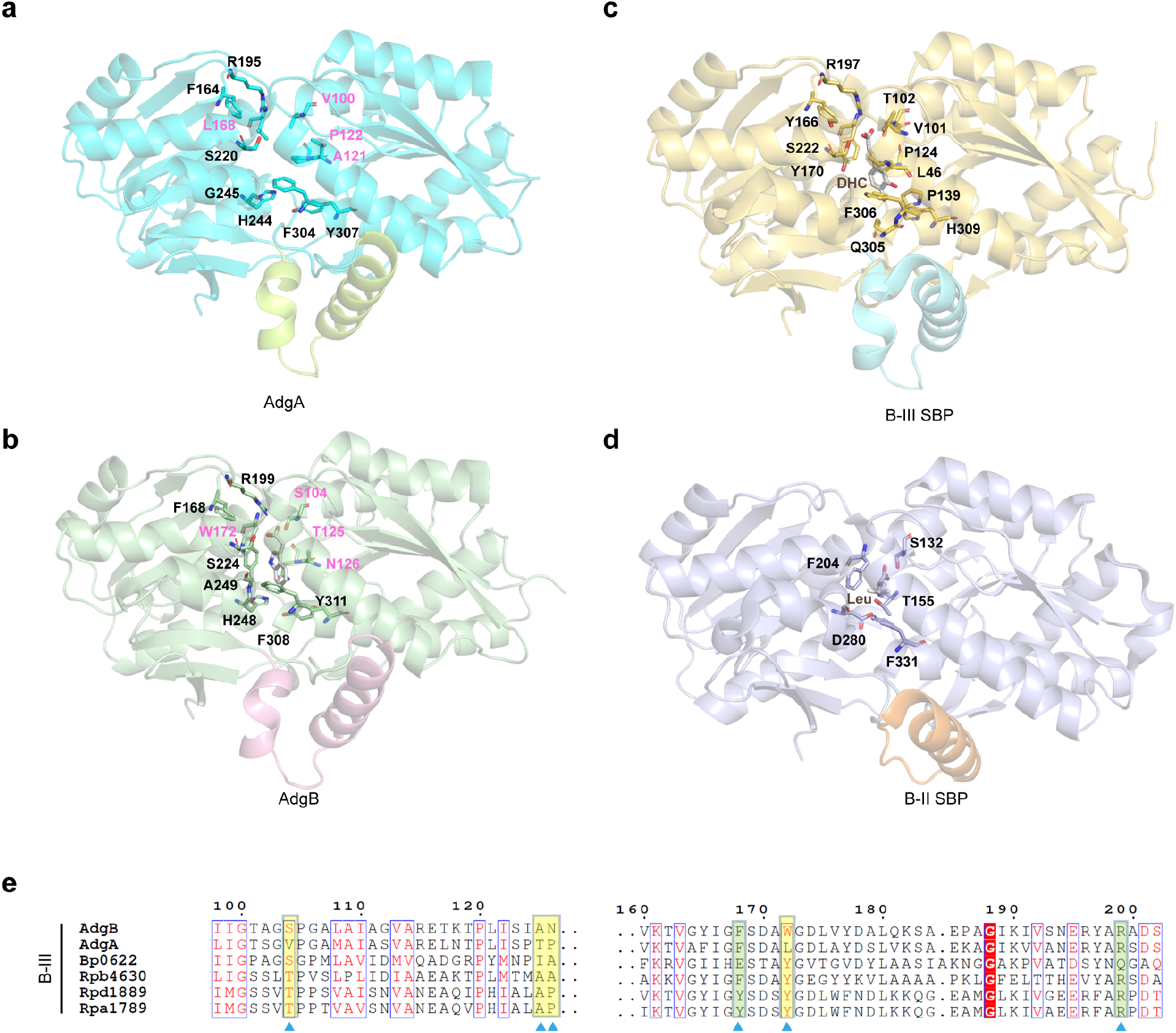
Comparison of AdgA, AdgB and other SBPs. **a**, Structure of AdgA predicted using AlphaFold2. The L2 linker is colored in green and the rest is in cyan. **b**, The calculated structure of AdgB complexed with IAA. Key residues interacting with IAA are shown in sticks representation. **c**, Homolog structure of AdgB belong to the B-III subcluster of SBPs. The structure of a SBP from *Rhodopseudomonas palustris* in complex with caffeic acid (DHC) is displayed (PDB: 4FB4). Key residues involved in substrate binding are shown. **d**, Structure of a B-II SBP. For comparison, the structure of a B-II type SBP in complex with a Leu amino acid substrate is displayed (PDB: 4GNR). **e**, Sequence alignment of AdgA, AdgB and B-III SBPs.

**Supplementary Figure 3.**
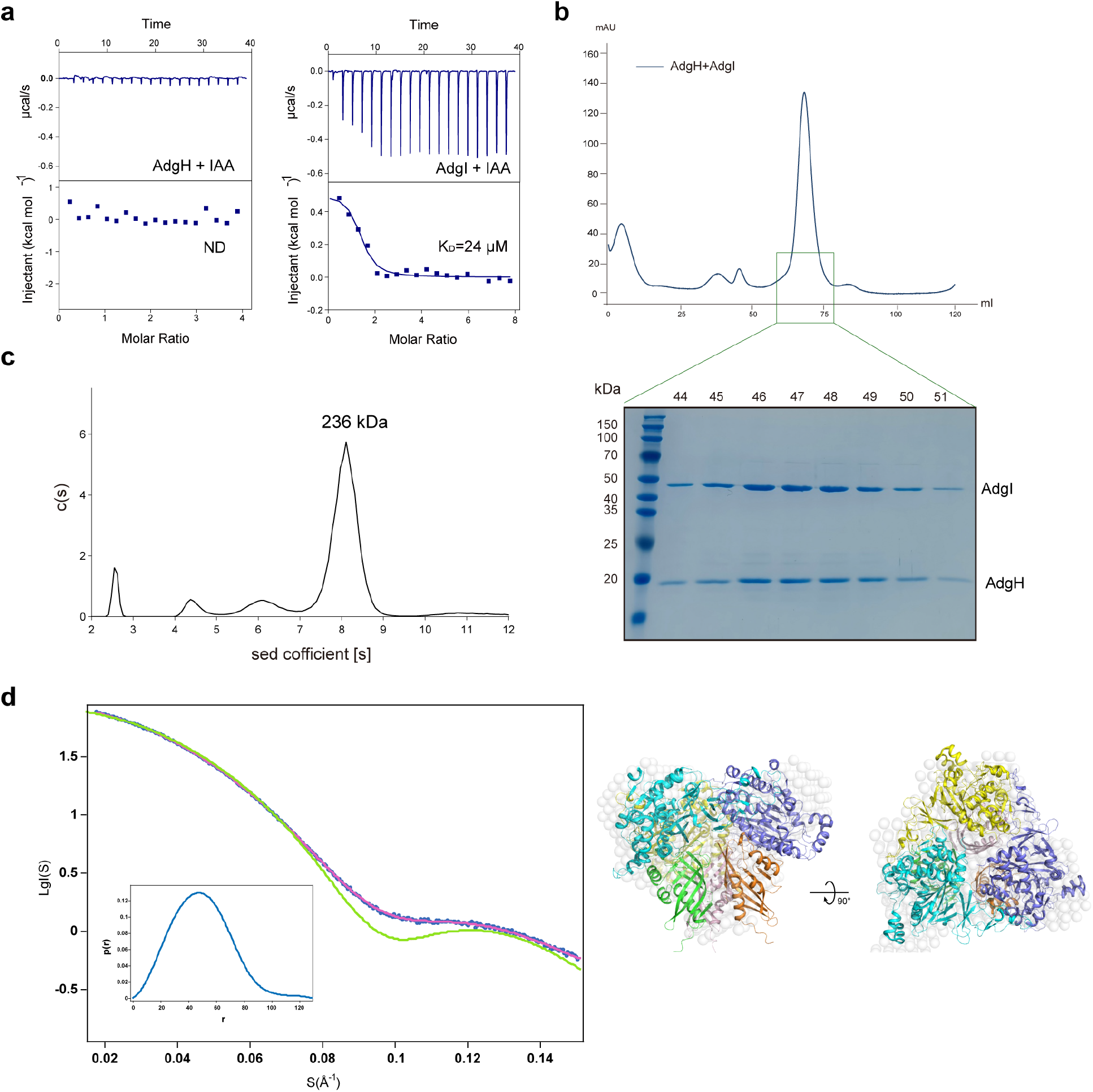
Biochemical and structural characterization of AdgI/AdgH. **a**, ITC measurements of IAA to AdgH or AdgI protein alone. ND: no binding detected. **b**, Size-exclusion chromatograms of AdgH and AdgI. The elution fractions from the complex peak were analyzed by SDS-PAGE gel (lower panel). **c**, Sedimentation coefficient distributions of AdgI /AdgH complex. **d**, Left panel: overlaid scattering pattern of the SAXS ab initio model (pink line) and the theoretical scattering profile of AdgI/AdgH complex structure (green line) with SAXS scattering data of AdgI/AdgH dioxygenase (blue dashed line). The inserted panel displays the P(r) distance distributions. Right: Superposition of the ab initio SAXS model (shown as white sphere) and cryo-EM structure of AdgI /AdgH complex.

**Supplementary Figure 4.**
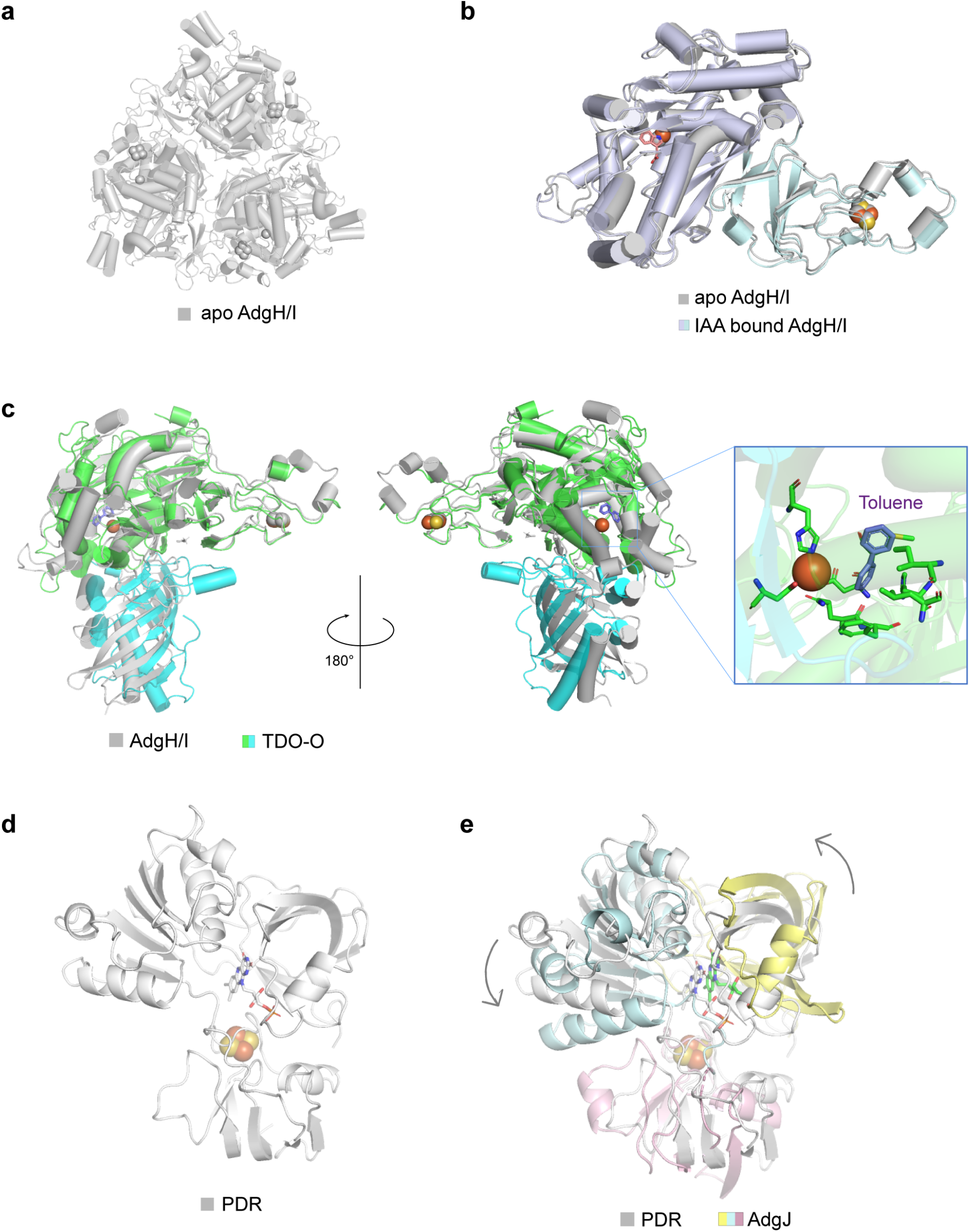
Structures of AdgI/AdgH dioxygenase and AdgJ reductase. **a**, AdgI/AdgH hetero hexamer in the unit cell of the apo crystallographic structure. **b**, Structural comparison of the apo and IAA-bound AdgI structures. The apo structure is colored in white and IAA-bound structure is colored by domains. **c**, Structural comparison of AdgI/AdgH-IAA with the homolog structure. For clarity, a single AdgI/AdgH heterodimer is displayed. Detailed insight into the homolog substrate binding pocket is shown (right panel). **d**, The crystal structure of PDR (PDB: 2PIA). **e**, Structural comparison of AdgJ with PDR. PDR is colored in white and AdgJ is colored by domains. The arrows indicate the relative domain rotations of AdgJ.

**Supplementary Figure 5.**
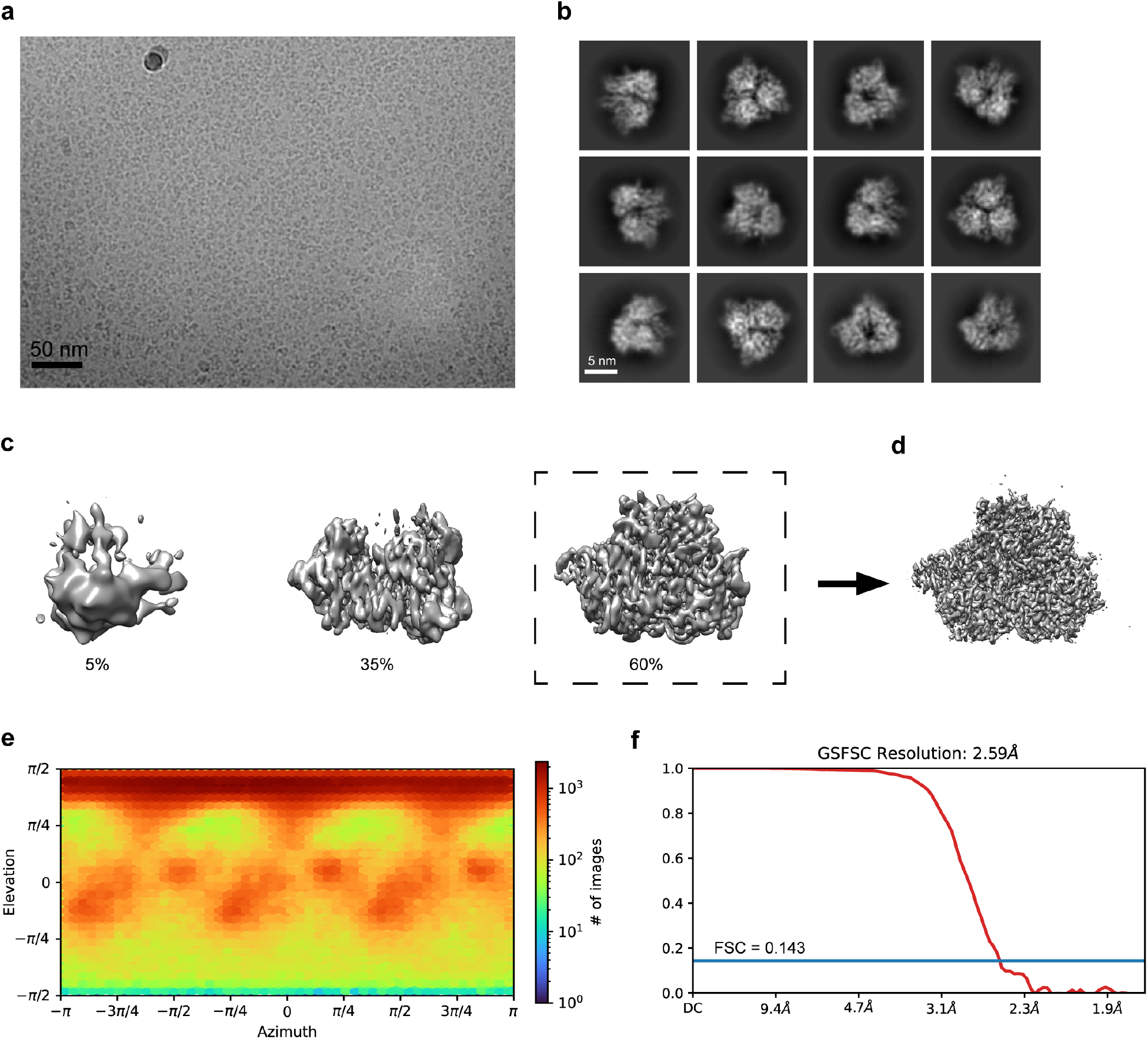
Cryo-EM data processing for the AdgI/AdgH-IAA complex. **a**, A representative raw cryo-EM micrograph. Scale bar: 50 nm. **b**, Representative 2D class averages. **c**, 3D classification. Class 1 represent the bad reconstruction. Class 2 represents the broken protein complex. Class 3 represents the intact protein complex. **d**, The final cryo-EM map of 3D refinement on Class 3. **e**, Viewing distribution of the 3D refinement. **f**, The Fourier shell correlation (FSC) curve of the reconstruction. The 0.143 gold standard FSC cutoff was used to determine the final resolution.

**Supplementary Figure 6.**
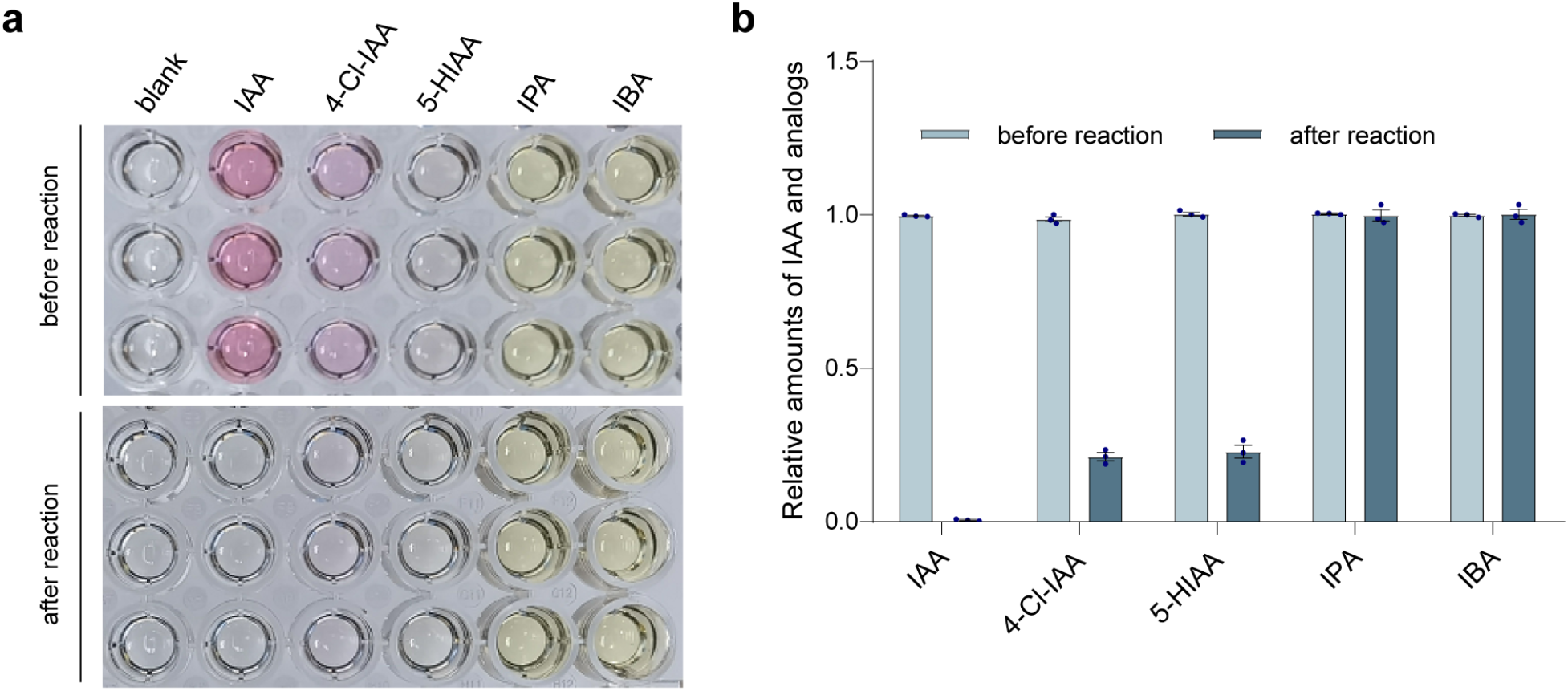
Degradataion of IAA and analogs by AdgI/AdgH and AdgJ. **a**, Color development of IAA and IAA analogs with the Salkowski reagent before and after treatments with AdgI/AdgH and AdgJ. **b**, Measurement of the relative amounts of IAA or IAA analogs after the oxidation reaction.

**Supplementary Figure 7.**
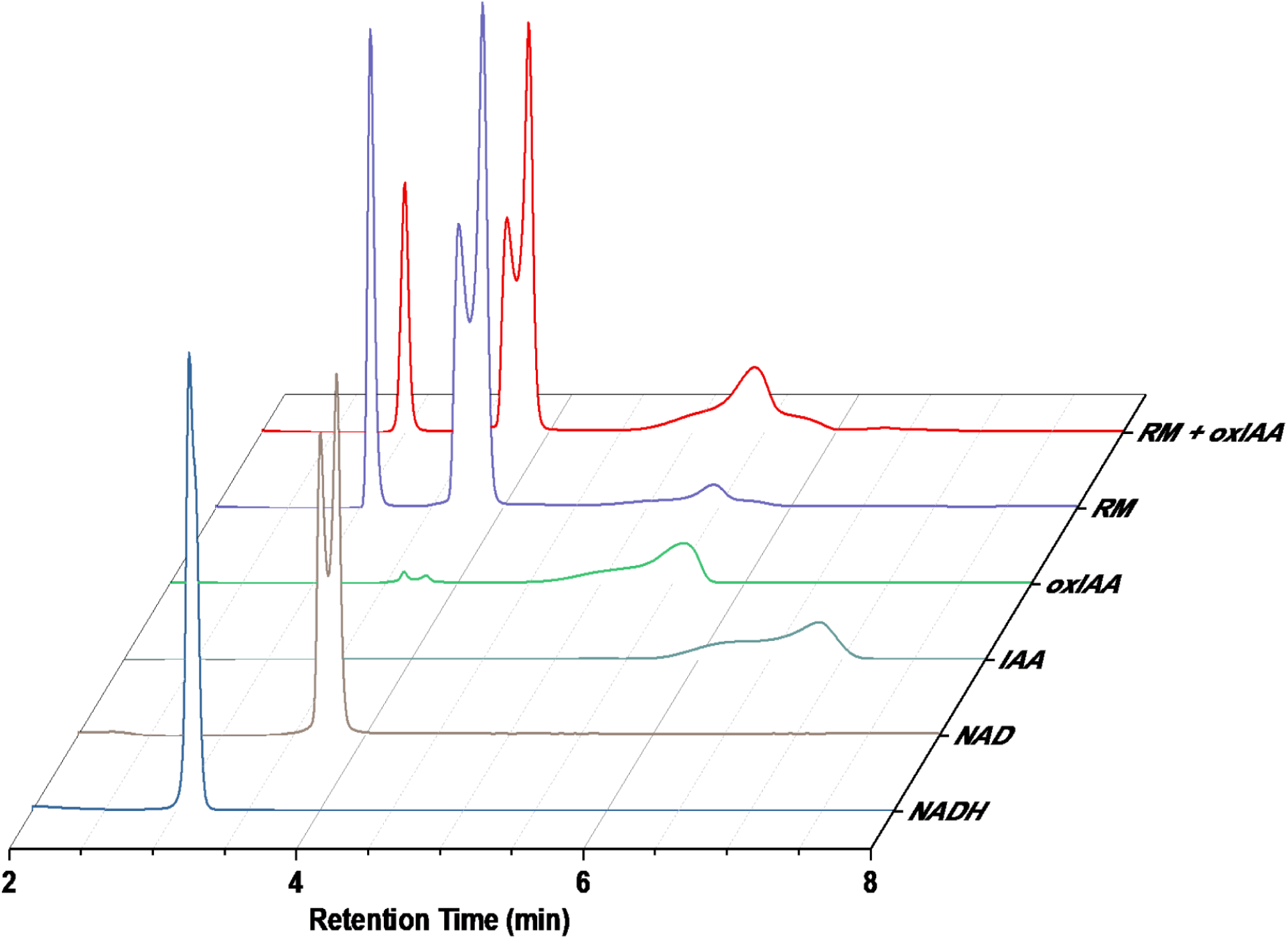
IAA metabolite analysis by HPLC. The reacted mixture (RM) was analyzed with HPLC. Analyses for NADH, NAD, IAA and oxIAA were also performed to assign the peaks.

**Supplementary Figure 8.**
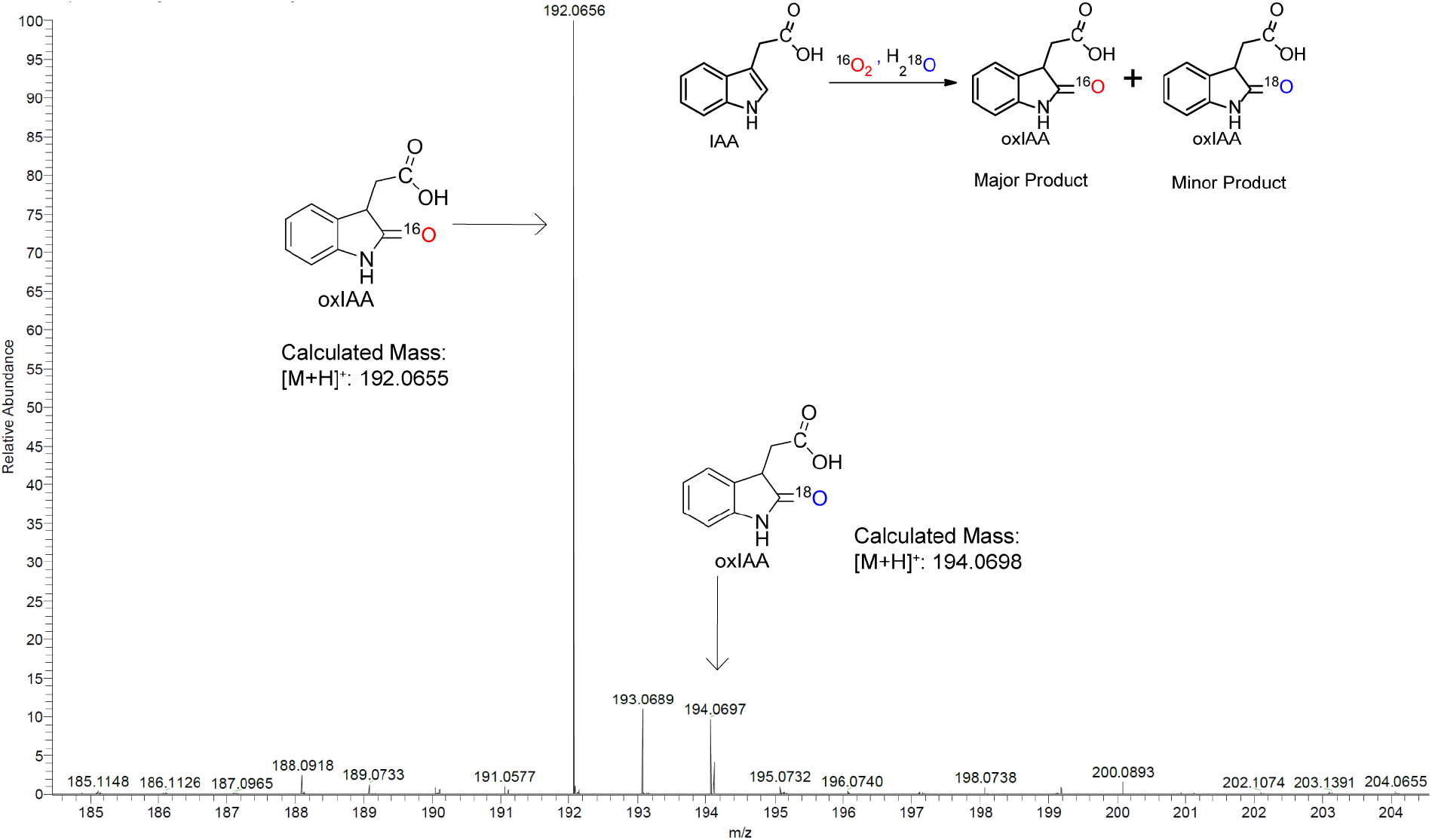
HRMS spectrum of isolated degradation product (oxIAA) of IAA from the isotopic experiment, which was carried out in the solution containing 95% H_2_O^18^ (v/v).

## Notes

### Competing Interest Statement

The authors have declared no competing interest.

## References

1 Brumos, J. et al. Local Auxin Biosynthesis Is a Key Regulator of Plant Development. Dev Cell 47, 306–318 e305, doi:10.1016/j.devcel.2018.09.022 (2018).

2 Grieneisen, V. A., Xu, J., Maree, A. F., Hogeweg, P. & Scheres, B. Auxin transport is sufficient to generate a maximum and gradient guiding root growth. Nature 449, 1008–1013, doi:10.1038/nature06215 (2007).

3 Hayashi, K. I. et al. The main oxidative inactivation pathway of the plant hormone auxin. Nat Commun 12, 6752, doi:10.1038/s41467-021-27020-1 (2021).

4 Duca, D., Lorv, J., Patten, C. L., Rose, D. & Glick, B. R. Indole-3-acetic acid in plant-microbe interactions. Antonie Van Leeuwenhoek 106, 85–125, doi:10.1007/s10482-013-0095-y (2014).

5 Ludwig-Muller, J. Bacteria and fungi controlling plant growth by manipulating auxin: balance between development and defense. J Plant Physiol 172, 4–12, doi:10.1016/j.jplph.2014.01.002 (2015).

6 Van Puyvelde, S. et al. Transcriptome analysis of the rhizosphere bacterium Azospirillum brasilense reveals an extensive auxin response. Microb Ecol 61, 723–728, doi:10.1007/s00248-011-9819-6 (2011).

7 Duran, P. et al. Microbial Interkingdom Interactions in Roots Promote Arabidopsis Survival. Cell 175, 973–983 e914, doi:10.1016/j.cell.2018.10.020 (2018).

8 Leveau, J. H. & Gerards, S. Discovery of a bacterial gene cluster for catabolism of the plant hormone indole 3-acetic acid. FEMS Microbiol Ecol 65, 238–250, doi:10.1111/j.1574-6941.2008.00436.x (2008).

9 Laird, T. S., Flores, N. & Leveau, J. H. J. Bacterial catabolism of indole-3-acetic acid. Appl Microbiol Biotechnol 104, 9535–9550, doi:10.1007/s00253-020-10938-9 (2020).

10 Scott, J. C., Greenhut, I. V. & Leveau, J. H. Functional characterization of the bacterial iac genes for degradation of the plant hormone indole-3-acetic acid. J Chem Ecol 39, 942–951, doi:10.1007/s10886-013-0324-x (2013).

11 Sadauskas, M., Statkeviciute, R., Vaitekunas, J. & Meskys, R. Bioconversion of Biologically Active Indole Derivatives with Indole-3-Acetic Acid-Degrading Enzymes from Caballeronia glathei DSM50014. Biomolecules 10, doi:10.3390/biom10040663 (2020).

12 Sun, S. L. et al. The Plant Growth-Promoting Rhizobacterium Variovorax boronicumulans CGMCC 4969 Regulates the Level of Indole-3-Acetic Acid Synthesized from Indole-3-Acetonitrile. Appl Environ Microbiol 84, doi:10.1128/AEM.00298-18 (2018).

13 Finkel, O. M. et al. A single bacterial genus maintains root growth in a complex microbiome. Nature 587, 103–108, doi:10.1038/s41586-020-2778-7 (2020).

14 Berntsson, R. P., Smits, S. H., Schmitt, L., Slotboom, D. J. & Poolman, B. A structural classification of substrate-binding proteins. FEBS Lett 584, 2606–2617, doi:10.1016/j.febslet.2010.04.043 (2010).

15 Scheepers, G. H., Lycklama, A. N. J. A. & Poolman, B. An updated structural classification of substrate-binding proteins. FEBS Lett 590, 4393–4401, doi:10.1002/1873-3468.12445 (2016).

16 Petersson, S. V. et al. An auxin gradient and maximum in the Arabidopsis root apex shown by high-resolution cell-specific analysis of IAA distribution and synthesis. Plant Cell 21, 1659–1668, doi:10.1105/tpc.109.066480 (2009).

17 Greenhut, I. V., Slezak, B. L. & Leveau, J. H. J. iac Gene Expression in the Indole-3-Acetic Acid-Degrading Soil Bacterium Enterobacter soli LF7. Appl Environ Microbiol 84, doi:10.1128/AEM.01057-18 (2018).

18 Brandl, M. T. & Lindow, S. E. Contribution of indole-3-acetic acid production to the epiphytic fitness of erwinia herbicola. Appl Environ Microbiol 64, 3256–3263, doi:10.1128/AEM.64.9.3256-3263.1998 (1998).

19 Tan, K. et al. Structural and functional characterization of solute binding proteins for aromatic compounds derived from lignin: p-coumaric acid and related aromatic acids. Proteins 81, 1709–1726, doi:10.1002/prot.24305 (2013).

20 Chandravanshi, M., Samanta, R. & Kanaujia, S. P. Structural and thermodynamic insights into the novel dinucleotide-binding protein of ABC transporter unveils its moonlighting function. FEBS J 288, 4614–4636, doi:10.1111/febs.15774 (2021).

21 Jumper, J. et al. Highly accurate protein structure prediction with AlphaFold. Nature 596, 583–589, doi:10.1038/s41586-021-03819-2 (2021).

22 Butler, C. S. & Mason, J. R. Structure-function analysis of the bacterial aromatic ring-hydroxylating dioxygenases. Adv Microb Physiol 38, 47–84, doi:10.1016/s0065-2911(08)60155-1 (1997).

23 Dhindwal, S. et al. Structural Basis of the Enhanced Pollutant-Degrading Capabilities of an Engineered Biphenyl Dioxygenase. J Bacteriol 198, 1499–1512, doi:10.1128/JB.00952-15 (2016).

24 Ferraro, D. J., Gakhar, L. & Ramaswamy, S. Rieske business: structure-function of Rieske non-heme oxygenases. Biochem Biophys Res Commun 338, 175–190, doi:10.1016/j.bbrc.2005.08.222 (2005).

25 Correll, C. C., Batie, C. J., Ballou, D. P. & Ludwig, M. L. Phthalate dioxygenase reductase: a modular structure for electron transfer from pyridine nucleotides to [2Fe-2S]. Science 258, 1604–1610, doi:10.1126/science.1280857 (1992).

26 Gassner, G. T., Ludwig, M. L., Gatti, D. L., Correll, C. C. & Ballou, D. P. Structure and mechanism of the iron-sulfur flavoprotein phthalate dioxygenase reductase. FASEB J 9, 1411–1418, doi:10.1096/fasebj.9.14.7589982 (1995).

27 Szkop, M., Sikora, P. & Orzechowski, S. A novel, simple, and sensitive colorimetric method to determine aromatic amino acid aminotransferase activity using the Salkowski reagent. Folia Microbiol (Praha) 57, 1–4, doi:10.1007/s12223-011-0089-y (2012).

28 Ashikawa, Y. et al. Electron transfer complex formation between oxygenase and ferredoxin components in Rieske nonheme iron oxygenase system. Structure 14, 1779–1789, doi:10.1016/j.str.2006.10.004 (2006).

29 Conway, J. M. et al. Diverse MarR bacterial regulators of auxin catabolism in the plant microbiome. Nat Microbiol 7, 1817–1833, doi:10.1038/s41564-022-01244-3 (2022).

30 Maqbool, A. et al. The substrate-binding protein in bacterial ABC transporters: dissecting roles in the evolution of substrate specificity. Biochem Soc Trans 43, 1011–1017, doi:10.1042/BST20150135 (2015).

31 Fitzpatrick, C. R. et al. Assembly and ecological function of the root microbiome across angiosperm plant species. Proc Natl Acad Sci U S A 115, E1157–E1165, doi:10.1073/pnas.1717617115 (2018).

32 Thiergart, T. et al. Root microbiota assembly and adaptive differentiation among European Arabidopsis populations. Nat Ecol Evol 4, 122–131, doi:10.1038/s41559-019-1063-3 (2020).

33 Schuck, P. On the analysis of protein self-association by sedimentation velocity analytical ultracentrifugation. Anal Biochem 320, 104–124, doi:10.1016/s0003-2697(03)00289-6 (2003).

34 Schuck, P., Perugini, M. A., Gonzales, N. R., Howlett, G. J. & Schubert, D. Size-distribution analysis of proteins by analytical ultracentrifugation: strategies and application to model systems. Biophys J 82, 1096–1111, doi:10.1016/S0006-3495(02)75469-6 (2002).

35 Petoukhov, M. V. et al. New developments in the ATSAS program package for small-angle scattering data analysis. J Appl Crystallogr 45, 342–350, doi:10.1107/S0021889812007662 (2012).

36 Manalastas-Cantos, K. et al. ATSAS 3.0: expanded functionality and new tools for small-angle scattering data analysis. J Appl Crystallogr 54, 343–355, doi:10.1107/S1600576720013412 (2021).

37 Konarev, P. V., Volkov, V. V., Sokolova, A. V., Koch, M. H. J. & Svergun, D. I. PRIMUS: a Windows PC-based system for small-angle scattering data analysis. Journal of Applied Crystallography 36, 1277–1282, doi:doi:10.1107/S0021889803012779 (2003).

38 Svergun, D. Determination of the regularization parameter in indirect-transform methods using perceptual criteria. Journal of Applied Crystallography 25, 495–503, doi:doi:10.1107/S0021889892001663 (1992).

39 Svergun, D. I. Restoring low resolution structure of biological macromolecules from solution scattering using simulated annealing. Biophys J 76, 2879–2886, doi:10.1016/S0006-3495(99)77443-6 (1999).

40 Potterton, L. et al. CCP4i2: the new graphical user interface to the CCP4 program suite. Acta Crystallogr D Struct Biol 74, 68–84, doi:10.1107/S2059798317016035 (2018).

41 Emsley, P., Lohkamp, B., Scott, W. G. & Cowtan, K. Features and development of Coot. Acta Crystallogr D Biol Crystallogr 66, 486–501, doi:10.1107/S0907444910007493 (2010).

42 Adams, P. D. et al. PHENIX: building new software for automated crystallographic structure determination. Acta Crystallogr D Biol Crystallogr 58, 1948–1954, doi:10.1107/s0907444902016657 (2002).

43 Afonine, P. V. et al. Towards automated crystallographic structure refinement with phenix.refine. Acta Crystallogr D Biol Crystallogr 68, 352–367, doi:10.1107/S0907444912001308 (2012).

44 Trott, O. & Olson, A. J. AutoDock Vina: improving the speed and accuracy of docking with a new scoring function, efficient optimization, and multithreading. J Comput Chem 31, 455–461, doi:10.1002/jcc.21334 (2010).

45 Eastman, P. et al. OpenMM 7: Rapid development of high performance algorithms for molecular dynamics. PLoS Comput Biol 13, e1005659, doi:10.1371/journal.pcbi.1005659 (2017).

46 Roe, D. R. & Cheatham, T. E., 3rd. PTRAJ and CPPTRAJ: Software for Processing and Analysis of Molecular Dynamics Trajectory Data. J Chem Theory Comput 9, 3084–3095, doi:10.1021/ct400341p (2013).

47 Maier, J. A. et al. ff14SB: Improving the Accuracy of Protein Side Chain and Backbone Parameters from ff99SB. J Chem Theory Comput 11, 3696–3713, doi:10.1021/acs.jctc.5b00255 (2015).

48 Zivanov, J. et al. New tools for automated high-resolution cryo-EM structure determination in RELION-3. Elife 7, doi:10.7554/eLife.42166 (2018).

49 Zheng, S. Q. et al. MotionCor2: anisotropic correction of beam-induced motion for improved cryo-electron microscopy. Nat Methods 14, 331–332, doi:10.1038/nmeth.4193 (2017).

50 Zhang, K. Gctf: Real-time CTF determination and correction. J Struct Biol 193, 1–12, doi:10.1016/j.jsb.2015.11.003 (2016).

51 Punjani, A., Rubinstein, J. L., Fleet, D. J. & Brubaker, M. A. cryoSPARC: algorithms for rapid unsupervised cryo-EM structure determination. Nat Methods 14, 290–296, doi:10.1038/nmeth.4169 (2017).

52 Liebschner, D. et al. Macromolecular structure determination using X-rays, neutrons and electrons: recent developments in Phenix. Acta Crystallogr D Struct Biol 75, 861–877, doi:10.1107/S2059798319011471 (2019).

53 Afonine, P. V. et al. New tools for the analysis and validation of cryo-EM maps and atomic models. Acta Crystallogr D Struct Biol 74, 814–840, doi:10.1107/S2059798318009324 (2018).

54 Adams, P. D. et al. PHENIX: a comprehensive Python-based system for macromolecular structure solution. Acta Crystallogr D Biol Crystallogr 66, 213–221, doi:10.1107/S0907444909052925 (2010).

55 Williams, C. J. et al. MolProbity: More and better reference data for improved all-atom structure validation. Protein Sci 27, 293–315, doi:10.1002/pro.3330 (2018).

